# Histone hyperacetylation-linked upregulation of KRAB zinc finger proteins impedes glial differentiation in Huntington’s disease

**DOI:** 10.64898/2025.11.30.691437

**Authors:** Peter Brøgger, John N. Mariani, Dennis Salinas, Benjamin Mansky, Steven J. Schanz, Nguyen P. T. Huynh, Rossana Foti, Steven A. Goldman

**Affiliations:** Center for Translational Neuromedicine, University of Copenhagen Faculty of Health, Copenhagen 2200, Denmark; Center for Translational Neuromedicine, University of Rochester Medical Center, Rochester, NY 14642

**Keywords:** Glial progenitor cell, astrocyte, oligodendrocyte, pluripotent stem cells, RNA-Seq, ATAC-Seq, epigenetics, gene regulatory networks, glial differentiation, Huntington’s disease, zinc finger repressors, KRAB domains

## Abstract

Glial differentiation is impaired in Huntington disease (HD), contributing to both the synaptic dysfunction and hypomyelination of HD. Through combined epigenomic and transcriptomic profiling, we found that glial progenitor cells (hGPCs) generated from HD-derived human embryonic stem cells exhibit persistent histone hyperacetylation, enabling the ectopic expression of a broad set of KRAB zinc finger protein (KZFP) transcriptional repressors. Single-cell RNA-Seq analysis of HD hGPCs revealed that their aberrant KZFP expression was attended by the persistent expression of neural progenitor-stage genes relative to wild-type hGPCs. The HD hGPCs over-expressed the MYST family histone acetyltransferase KAT6B, which led to their hyperacetylation at H3K9 and associated DNA demethylation, and displayed abnormally open chromatin, particularly at promoters of chromosome 19 KZFP gene clusters. Among those KZFPs most differentially activated in HD hGPCs was the primate-specific *ZNF98*, whose overexpression in wild-type hGPCs recapitulated the HD-associated suppression of glial development. These data implicate abnormal histone hyperacetylation in HD glial progenitor cells, and its associated over-expression of recently evolved KZFP transcriptional repressors, as a critical mechanism by which both astrocytic and oligodendrocytic differentiation are impaired in HD.

## INTRODUCTION

Huntington’s disease (HD) is a monogenic neurodegenerative disorder characterized by progressive motor, cognitive, and behavioral deficits. It is caused by unstable CAG trinucleotide expansions in the first exon of the *Huntingtin* (*HTT*) gene, resulting in abnormally extended polyglutamine (polyQ) tract. HD is fully penetrant in individuals with ≥40 CAG repeats, with longer repeats correlating with earlier onset and more rapid disease progression^1,2^. While neurodegeneration is most prominent in the striatum due to the loss of selectively vulnerable medium spiny neurons (MSNs)^3^, both human and animal studies have recently highlighted dysfunctional oligodendrocytes and astrocytes in HD, likely subserving the white matter involution and circuit dysregulation observed in premanifest disease^4-9^.

We previously reported that mutant HTT-expressing human glial progenitor cells (hGPCs) – bipotential for oligodendrocytes and astrocytes, and produced from embryonic stem cells (hESCs) derived from HD blastocysts - recapitulated salient aspects of HD pathology when transplanted into mouse striata. These cells manifested impaired development in vivo compared to hGPCs derived from healthy wild-type siblings, with delays in both oligodendrocytic and astrocytic maturation^10,11^. In particular, HD hGPC-derived oligodendrocytes exhibited deficient myelination, while HD astrocytes were morphologically abnormal, paralleling aspects of glial pathology noted in HD animal models^12-14^. Transcriptional profiling of HD hGPCs revealed their downregulation of genes involved in myelination, axon guidance, synaptogenesis, and synaptic signaling^10^, along with the suppression of key drivers of glial differentiation including *NKX2.2, OLIG2, SOX10,* and *MYRF*, highlighting the cell-intrinsic nature of the glial differentiation defect in HD. Defective TCF7L2 signaling in particular has been implicated in the defective myelination program of R6/2 HD glial progenitors, which proved rescuable by TCF7L2 overexpression^15^. Acting as a stage-specific switch for glial differentiation^16^, TCF7L2 may promote the maturation of astrocytes as well, affecting their spatial tiling, gap junction coupling, and synaptic regulation^14,17^. Together, these observations suggest that transcriptional dysregulation in HD hGPCs disrupts glial lineage progression, yielding functional defects in both oligodendrocytes and astrocytes. Remarkably though, little is known about the chromatin landscape in HD glia or their progenitors, even though significant epigenetic changes – including altered DNA methylation and histone acetylation – have been noted in HD models and patient samples alike^18-20^.

Here, we asked if the widespread suppression of glial differentiation factors in HD hGPCs is mediated by mHTT-dependent epigenomic alterations. We profiled multiple layers of epigenetic regulation, including chromatin accessibility via ATAC-seq, histone post-translational modifications using CUT&Tag, and CpG DNA methylation via Infinium array. This integrated analysis revealed significant chromatin relaxation in HD hGPCs, as marked by H3K9 hyperacetylation and DNA demethylation, at recently evolved KRAB zinc finger protein (KZFP) clusters on chromosome 19 (chr19), resulting in their promiscuous and heterochronic expression. While these KZFPs typically repress transposable elements (TEs), they also influence gene activity through both TE-embedded enhancers and LTR-derived promoters^21,22^. Notably, the number of upregulated KZFPs *increased* with CAG repeat length, in contrast to the CAG repeat length-dependent *repression* of glial differentiation factors^10^, suggesting a causal relationship between the CAG repeat length, the ectopic expression of KZFP repressors and impaired glial differentiation.

To investigate this association, we used single-cell RNA sequencing (scRNA-seq) of cultured HD hGPCs, paired with CUT&Tag assessment of histone acetylation marks, to reveal that HD hGPCs aberrantly overexpress the MYST family histone acetyltransferase (HAT) KAT6B (MORF), along with many KZFPs. Accordingly, KAT6B sites were found to intersect with the aberrantly open CpG islands (CpGIs) at H3K9 hyperacetylated KZFP gene promoters across chr19.

Motif enrichment analysis among HD-associated, differentially expressed hGPC genes identified ZNF98, a hominid-specific KZFP, as a candidate disruptor of glial differentiation. Lentiviral overexpression of ZNF98 in healthy hGPCs recapitulated salient features of the transcriptional dysregulation of HD hGPCs, including suppression of critical regulators of contact-mediated functions, such as netrin surface receptors, and increased expression of progenitor-associated genes, such as GSX2 and CD133. Together, these data indicate that the mHTT-linked overactivity of KAT6B and the consequent H3K9 hyperacetylation in HD hGPCs promotes their aberrant expression of KZFP repressors. These KZFPs in turn repress glial maturation-associated transcriptional programs, thus hindering both astrocytic and oligodendrocytic maturation. Among the many KZFPs aberrantly expressed in HD hGPCs, the hominid-specific ZNF98 reproduces salient aspects of the HD glial phenotype when expressed in healthy sibling glia.

## RESULTS

### HD hGPCs manifest widespread promoter histone hyperacetylation

We performed cleavage under targets and tagmentation (CUT&Tag) to profile histone modifications in hGPCs derived from the female human embryonic stem cell (hESC) lines GENEA19 (G19, WT) and GENEA20 (G20, HD) (n=4/group). All cells were studied at 160-180 days in vitro (DIV), corresponding to roughly 30-50 days after their in vitro acquisition of glial progenitor cell phenotype. The G20 line models late-onset HD (48CAG repeats in *mHTT*), and is a sibling to the healthy G19 control line (18/16 CAG) (**Figure 1A**). To explore the chromatin landscape regulating gene expression in HD hGPCs, we assessed the histone tail marks H3K4me3, H3K27ac, and H3K9ac – all associated with active transcription and open chromatin. Principal component analysis (PCA) demonstrated clear separation between WT and HD-derived hGPCs for H3K27ac and H3K9ac (**Figure S1A**), while the clustering of H3K4me3 samples was less distinct, suggesting that differential chromatin activation is mediated primarily through acetylation.

**Figure 1.**
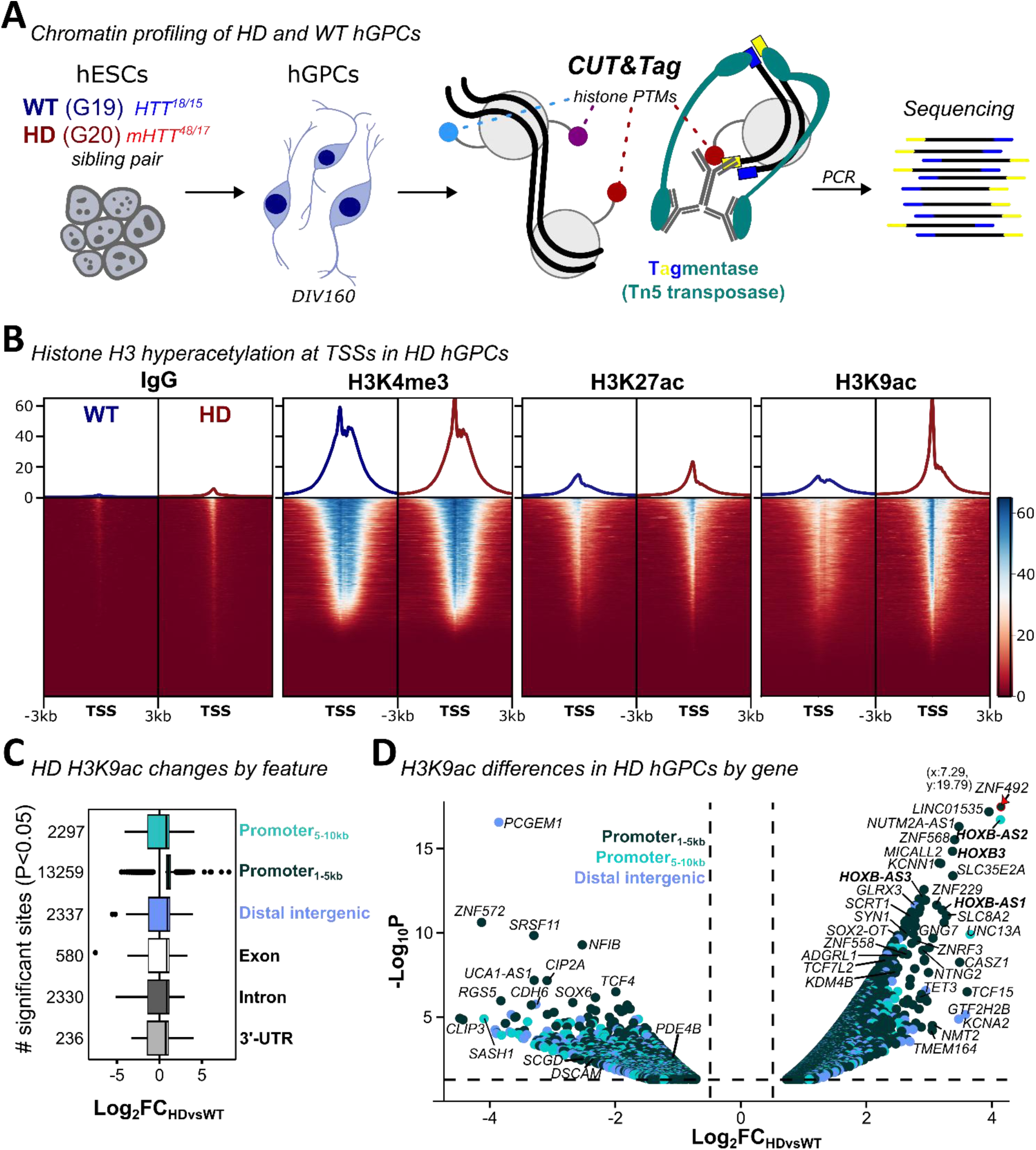
HD hGPCs exhibit profound histone H3 hyperacetylation at gene promoters. **A.** Human glial progenitor cells (hGPCs) differentiated from the sibling embryonic stem cell lines G19 (WT) and G20 (HD, n=4/group) were processed for cleavage under targets and tagmentation (CUT&Tag) following 160 days in vitro (DIV160) for characterization of histone tail post translational modifications (PTMs) H3K4me3, H3K27ac, and H3K9ac associated with gene activation. **B.** Profile plots show the median signal intensities over refseq transcription start sites (TSSs) of H3K4me3, H3K27ac, H3K9ac, and the IgG negative control, with HD hGPCS (red) exhibiting marked H3K9 hyperacetylation in comparison to healthy WT (blue) hGPCs. **C.** Sites showing significant H3K9ac differences identified by *DiffBind* (p<0.05) were most frequently found at promoter regions within 5kb of TSSs and showed H3K9ac accumulation in HD hGPCs **D.** Significantly different H3K9ac sites are shown for promoter-associated sites (teal colors) or distal intergenic elements (blue) following gene annotation. The most markedly different H3K9ac differences are indicated. For transcripts with multiple sites of significance, only the most significant were plotted. Stippled lines mark plotting thresholds p<0.05 and |Log_2_FC|>0.5. The red arrow indicates an outlier, which was lowered to fit the plot area (plot coordinates are shown).

HD hGPCs exhibited pronounced increases in H3K27ac and especially H3K9ac at gene transcription start sites (TSSs), predicting their differentially active transcription relative to the same genes in WT hGPCs (**Figure 1B**). Of approximately 20,000 H3K9ac sites identified as significantly different by *DiffBind* (*p<0.05*), the majority represented increases at promoters within 10kb of TSSs (**Figure 1C**). Overall, 12,382 promoters showed differential H3K9ac accumulation (*Log_2_FC>0.5*) in HD hGPCs (**Figure 1D**). Especially strong H3K9ac gains were detected at the HoxB cluster across *HOXB2, -3,* and *-4,* and at promoters of synaptic genes including *SLC8A2, UNC13A, KCNN1, ADGRL1,* and *SYN1*; these were frequently accompanied by increases in H3K4me3 and H3K27ac (**Figures S1B**-**S1C**), suggesting the concurrent over-activation of these genes in HD hGPCs. Interestingly, aside from *SYN1,* these synaptic genes are located on chr19, a chromosome characterized by high gene density and its dynamic localization near transcriptionally active nuclear speckles^23,24^.

Conversely, relatively fewer promoters in HD hGPCs exhibited significant H3K9ac loss (3,174 sites), but these included genes related to astrocyte and oligodendrocyte differentiation such as *SOX6*^25^ and *SASH1*^26^, and were accompanied by the concomitant loss of the activating markers H3K4me3 and H3K27ac; this pattern was consistent with the downregulated transcripts identified in HD hGPCs (**Figures 1D** and **S1D**). Of note, despite the high similarity between H3K4me3 samples from G19 and G20 hGPCs, we found significant loss of H3K4me3 at the promoters of a number of downregulated glial genes (Log_2_FC<-0.5, p<0.05); these included *PCDH15, NKX2-2, SOX6, GPR17, EYA1, DOCK10, SASH1, and NCAM2* (**Figure S1E**), whose collective down-regulation both predicted and reflected the impaired glial development and functional deficits of HD hGPCs^10^.

### DNA demethylation marks ZFPs and WNT effectors for activation in HD hGPCs

DNA methylation at promoter CpG islands (CpGIs) is a stable mark associated with gene silencing, while demethylation following histone acetylation is required for sustained gene expression^27,28^. To compare genome-wide DNA methylation patterns between HD (G20) and sibling wild-type (WT; G19) hGPCs, we used Infinium 450k methylation arrays at DIV180 (n=4 replicates/line). Principal component analysis showed clear separation between the HD and WT samples (**Figure S2A**), identifying 44,084 significantly differentially methylated (DM) sites (*limma,* |Log_2_FC|>0.5, p<0.05). DM sites were categorized based on their overlap with CpGI regions (*islands, shores, shelves, open sea*, which were defined respectively by their distance to CpGIs; 0, 2, 4, and >4kb^29^), as well as by their genomic context (promoter and first exon sites, gene bodies, 3′-UTRs, or distal intergenic sites) (**Figure 2A**). Most DM sites mapped to promoters (27,577), followed by intragenic (9,378), intergenic (5,974), and downstream regions (1,155, **Figure S2B**). 16,136 of these DM sites were found in *open sea* regions, while many still mapped close to gene transcription start sites (TSSs) and were otherwise mainly associated with CpG *islands* (13,463 sites in total, **Figures S2B**-**S2C**) and *shores* (10,970).

**Figure 2.**
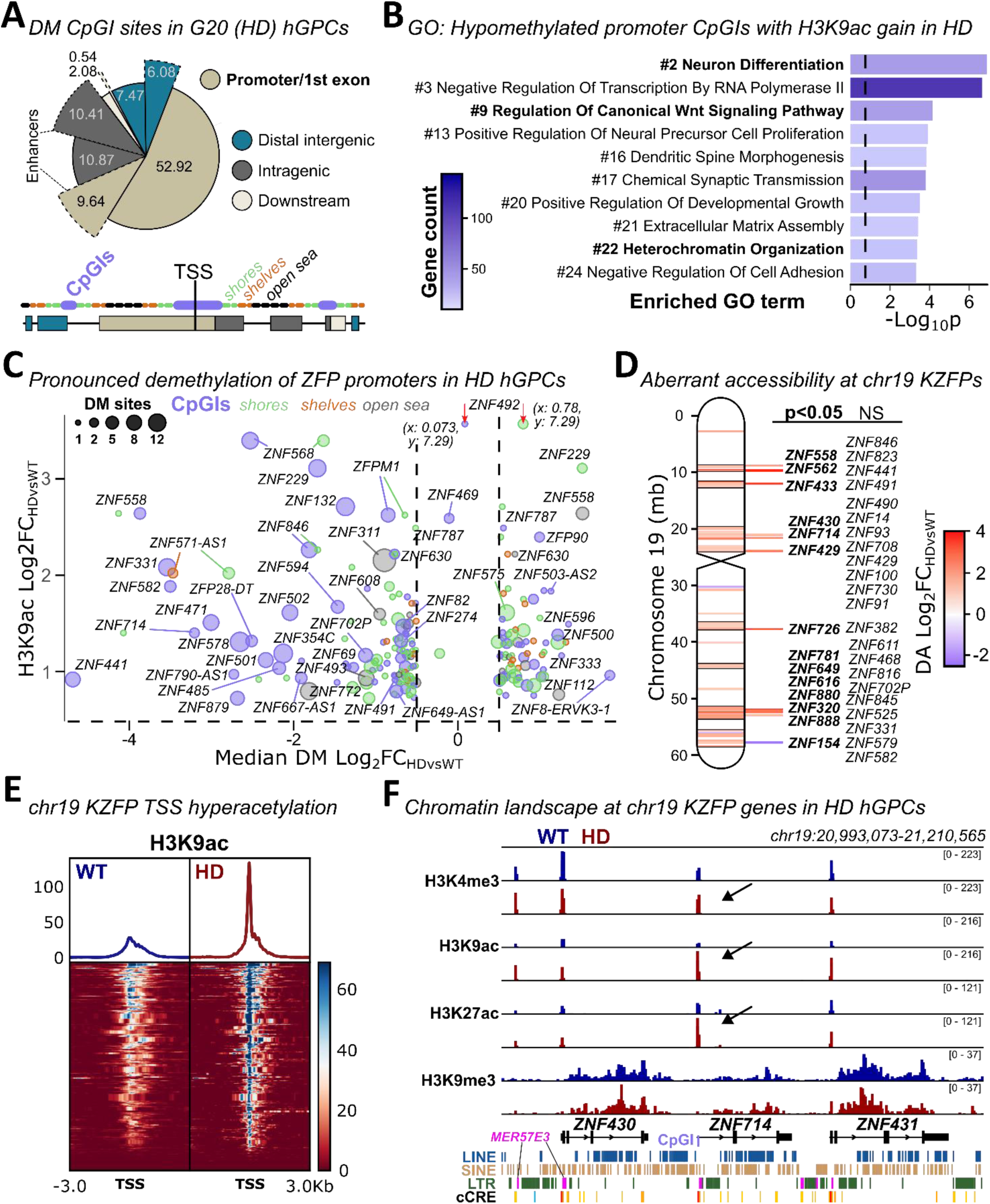
HD hGPCs exhibit aberrant accessibility at KZFP gene promoters on chromosome 19. **A.** The pie chart summarizes the annotated gene features of significantly differentially methylated (DM) sites identified between G20 (HD) and G19 (WT) hGPCs (n=4/group) using *limma*. The accompanying overview categorizes DM sites by gene feature type and by proximity to annotated CpG islands (CpGIs), with CpGIs, shores, shelves, and open sea corresponding to distances of 0, 2, 4, and >4kb from the nearest CpGI. **B.** The top enriched gene ontology (GO) terms for significantly demethylated CpGI sites (Log_2_FCs<-0.5 & p<0.05, *enrichR*) that overlap significantly H3K9 hyperacetylated promoters (Log_2_FCs>0.5 & p<0.05) are shown. These terms were picked from the 25 most significant categories (see **Table S1**). **C.** Pronounced DNA demethylation coincided with marked H3K9 hyperacetylation at several promoters of zinc finger protein (ZFP) genes, as illustrated in this bubble plot. The plot shows H3K9ac accumulation at these loci relative to the median Log_2_FC derived from multiple significant (p<0.05) DM sites, with bubble size reflecting the number sites per locus. Red arrows denote outliers adjusted to remain within the plotting range (coordinates indicated). **D.** Differential chromatin accessibility (ATAC-seq, *DiffBind*) in G20 (HD) versus G19 (WT) hGPCs (n=2/group) revealed a tendency toward aberrantly relaxed chromatin states at annotated ZFP gene promoters and distal intergenic regions across chromosome 19 (chr19). The most pronounced differences (|Log_2_FC|>1) are highlighted using *ggbio*, with significant changes (p<0.05) shown in bold. Many of these ZFP genes belong to the KRAB ZFP (KZFP) gene family, which is clustered across chr19 (see **Table S2**). **E.** Median H3K9ac signal intensities over the TSSs of these KZFP genes on chr19 confirm that these loci are highly acetylated in the G20 (HD) hGPCs. **F.** Representative *IGV* tracks illustrate KZFP genes from the chr19 p12 cluster displaying H3K9 hyperacetylation in G20 (HD) hGPCs accompanied by increased H3K4me3 and H3K27ac signals indicative of transcriptional activation (arrows). Little change was observed in the characteristic H3K9me3 heterochromatin (n=4), across the transposable element (TE)-rich KZFP regions.

Some HD hGPCs exhibited significant DNA hypomethylation of their CpGIs (Log_2_FC<-0.5, p<0.05) and overlapping H3K9-hyperacetylated promoters; these regions marked transcriptionally active genes in HD hGPCs. Gene ontology (GO) enrichment analysis (*enrichR*) of these genes revealed their overrepresentation of WNT signaling and neuronal differentiation-associated transcripts (**Figure 2B** and **Table S1**). These HD-associated, differentially-overexpressed genes included *WNT1, -2, -3, -5A, -5B, -7B, and -10B, AXIN2, CTNND2 APC2, FZD1* and *-8, RUNX2, LHX2, and GSX1* (**Figure S2D**).

Widespread transcriptional dysregulation in HD hGPCs was suggested by the top enriched GO terms related to transcription (**Table S1**). Of note, the genes encoding several zinc finger proteins (ZFPs) were identified in these categories, many displaying pronounced DNA hypomethylation at promoter CpGIs in hyperacetylated regions (**Figure 2C**), while smaller degrees of relative hypomethylation were observed at shores, shelves and open sea regions.

### Aberrantly open chromatin architecture at chr19 KZFP clusters in HD hGPCs

We next asked if HD hGPCs exhibited increased chromatin accessibility, relative to WT hGPCs, at those ZFP promoters marked for active transcription. To that end, we performed ATAC-seq (*assay for transposase-accessible chromatin sequencing*)^30^ on HD (G20) and WT (G19) hGPCs at DIV180 (n=2/group, **Figure S2E**). We found that the HD hGPCs indeed manifested increased chromatin accessibility at several clustered ZFP genes on chr19, specifically those of the Krüppel-associated box (KRAB) ZFP (KZFP) family (**Figures 2D** and **S2F**). In accord with their increased open chromatin and DNA demethylation, the chr19-cluster KZFP genes in HD hGPCs showed pronounced H3K9ac accumulation at their TSSs (**Figure 2E**). This was frequently accompanied by elevated H3K27ac and H3K4me3 activation signals (**Figure S2G**), particularly at *ZNF568, ZNF229, ZNF331, ZNF132, ZNF582, ZNF491,* and *ZNF714* with significantly demethylated promoter CpGIs (**Figures 2C** and **2F**), as well as at promoters in accessible chromatin regions, including *ZNF558, ZNF714, ZNF320, ZNF726, ZNF562,* and *ZNF429* (**Figure 2D**).

Of note, several of these KZFPs are evolutionarily recent in origin, and are restricted to primates, and in many cases even moreso, to hominids (see **Table S2**). Gene-spanning H3K9me3 heterochromatin associated with KZFPs (*atypical heterochromatin*)^31^ was studied as well, but appeared largely unaffected in HD (G20) hGPCs (**Figure 2F**). Together, these data highlight the selective increase in chromatin accessibility and active histone modifications at specific chr19 KZFP clusters in HD-derived glial progenitors, implicating aberrant epigenetic activation of these loci – a number of which are hominid-specific – in the pathogenesis of HD.

### HD hGPCs upregulate increasing chr19 KZFPs with mHTT CAG repeat length

To assess whether chromatin relaxation at chr19 KZFP clusters might correlate with the transcriptional patterns of HD hGPCs, we re-analyzed previously published bulk expression data^10^. This analysis revealed a progressive upregulation of the number of differentially expressed ZFP and KZFP genes with increasing CAG repeat length, across hGPCs derived from three different HD lines of 42, 44 and 48 CAGs (|Log_2_FC|>0.5, *p_adj_*<0.05; consistently identified by *DESeq2* and *edgeR*, **Figures 3A**-**3B** and **Table S3**). This relationship was reflected in the strong linear correlation between the number of dysregulated chr19 KZFP genes and CAG repeat length (**Figure 3C**).

**Figure 3.**
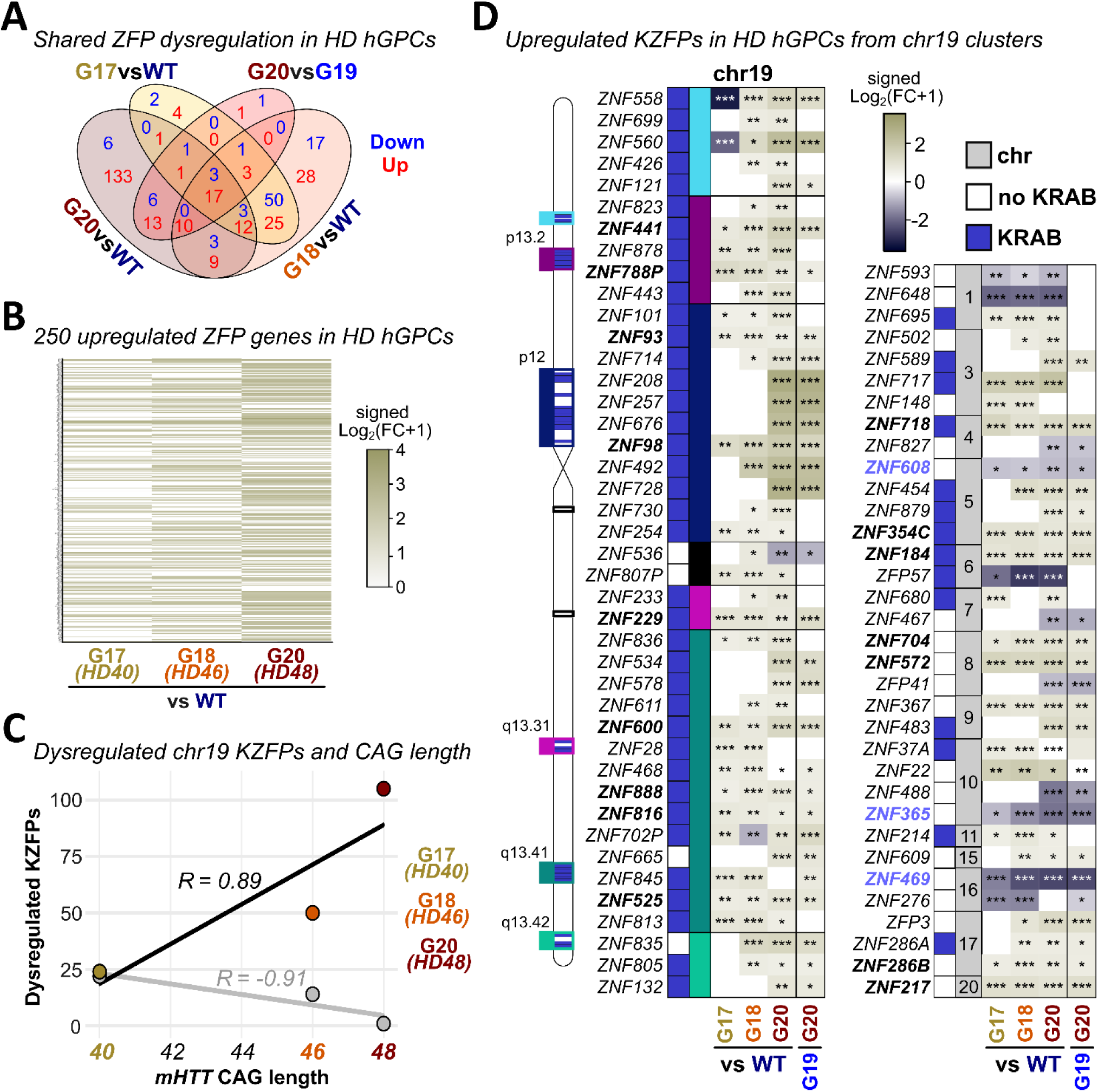
KZFP dysregulation in HD hGPCs increases with CAG repeat length. **A.** The Venn diagram depicts the numbers of significantly downregulated (blue) and upregulated (red) ZFP genes identified in three different independent HD hGPC lines (RNA-seq; |Log_2_FC|>0.5 & p_adj_<0.05; shared between *edgeR* and *DESeq2*; n=4-6/line; 3 HD lines and 2 WT lines^10^). All HD lines were compared relative to the pooled WT hGPC samples, with G20 additionally compared to its G19 sibling control line. **B.** Heatmap showing all significantly upregulated ZFPs (see **Table S3**) across HD hGPCs ordered by increasing mHTT CAG repeat lengths (*HD40-48* corresponding to lines G17, G18, and G20). Signed Log_2_FC transformation (sign(Log_2_FC)•Log2(|Log_2_FC|+1)) was applied to reduce outlier effects while preserving directionality. **C.** The number of significantly dysregulated KRAB ZFP (KZFP) genes expressed from chr19 increases linearly with CAG repeat length, as shown by Pearson correlation coefficients (R) for G17, G18 and G20 HD hGPCs. **D.** Heatmaps display significantly up- or downregulated ZFP genes on chr19 (left) and other chromosomes (right) in G20 (HD) hGPCs compared to its G19 (WT) sibling control, or differing significantly in at least two additional comparisons. Most chr19 ZFPs belong to the KZFP family (highlighted by blue boxes), with distinct KZFP clusters colored on the ideogram. Asterisks indicate significance; *P < 0.05; **P < 0.01; ***P < 0.001.

Key upregulated KZFP genes included *ZNF93, ZNF98*, *ZNF229, ZNF441, ZNF525, ZNF600, ZNF788P, ZNF816,* and *ZNF888,* which were elevated across all HD lines (**Figure 3D**). Other dysregulated KZFPs, such as *ZNF132* and *ZNF578*, were overly expressed specifically in G20 (HD) hGPCs, which displayed the most pronounced transcriptional dysregulation relative to WT hGPCs. The KZFP overactivity aligns well with the epigenetic changes observed in HD (G20) hGPCs, confirming that the mechanisms regulating coordinated KZFP expression are sensitive to the effects of mHTT. In contrast, downregulated ZFP genes predominantly belonged to KRAB-less transcription factor (TF) families, including *ZNF365* (also known as *DBZ*) – another regulator of oligodendrocyte differentiation^32^ – and *ZNF469* or *ZNF608*, whose roles in glial biology remain uncharacterized.

### Immature HD hGPCs exhibit heterochronic KZFP expression

To determine whether KZFP upregulation of HD hGPCs is widespread or restricted to specific cell stages, we performed 10X single-cell RNA sequencing (scRNA-seq) of WT and HD hGPCs (three independent lines/condition with HD lines carrying 40, 46 or 48 CAG repeats on one allele; see Methods). Cells were captured either directly at DIV160 (n=2/line), or after 19-20 weeks in vivo, following their transplantation into the corpus callosa (CC) of neonatal hypomyelinated *shiverer* x rag2^-/-^ (SR2) mice (**Figure 4A**, n=2–4 mice/line). Systematic assessment of glial surface marker expression (PDGFRA and CD44) in hGPCs by flow cytometry confirmed that these inductions were analogous (**Figure S3A**).

**Figure 4.**
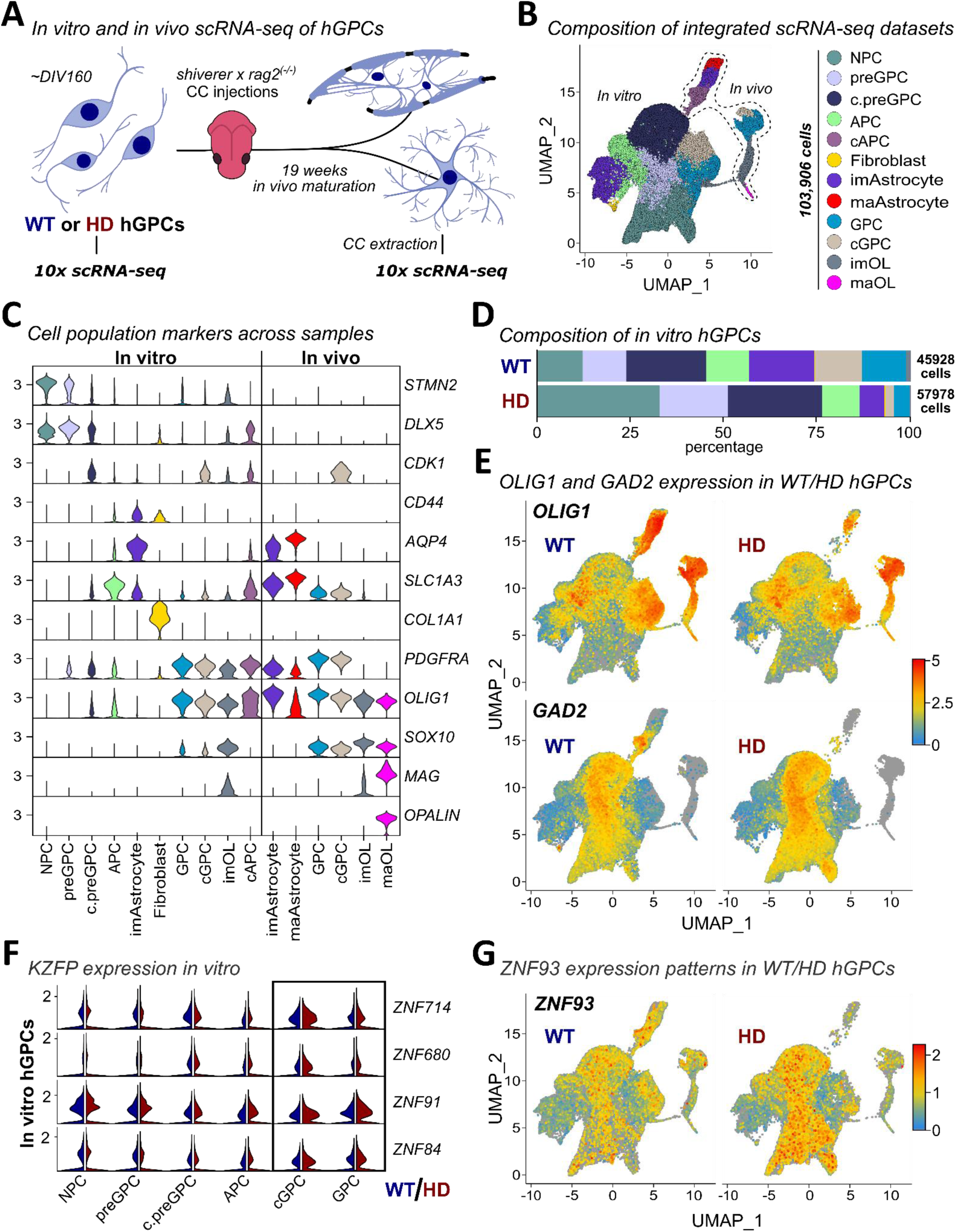
Single-cell profiling reveals altered maturation and persistent KZFP expression in HD hGPCs. **A.** WT and HD hGPCs (n=3 lines/condition) were cultured for 160 days in vitro (DIV160) and processed either directly single-cell RNA sequencing (scRNA-seq, 10X Genomics; n=2/line) or transplanted into the corpus callosum (CC) of neonatal SR2 pups (n=2-4/line). After 19 weeks, matured hGPCs were isolated and subjected to a second round of scRNA-seq. **B.** Uniform Manifold Approximation and Projection (UMAP) plots (*Seurat*) of all integrated datasets reveal distinct clustering of in vitro and in vivo-derived hGPCs with cell subtypes color-coded. **C.** Violin plots display the expression of canonical marker genes used to assign major cell identities. **D.** The composition of in vitro HD hGPCs suggest a shift toward immature populations compared with WT controls. **E.** Feature plots illustrate the expression of *GAD2 –* normally enriched in radial glia or neural progenitor cells (NPCs) but associated with GABAergic differentiation – as well as the gliogenic factor *OLIG1* across WT and HD hGPCs. **F.** Several KZFPs encoded on chr19, including *ZNF84, ZNF91, ZNF680*, and *ZNF714* show strong expression in the NPC-like subsets and remain upregulated in glia-leaning subpopulations of HD hGPCs. **G.** Expression feature plots highlight the high activity of the KZFP *ZNF93* within NPC clusters that are prevalent in HD in vitro samples.

After integration and clustering, the major cell types were identified based on canonical marker expression, showing very little overlap between the integrated in vitro and in vivo datasets (**Figures 4B**–4C and **Figure S3B**). These included cycling and non-cycling pre-glial progenitor cells, glial progenitor cells, and astrocyte precursor cells, as well as neural progenitor cells (clusters c.preGPC, preGPC, cGPC, GPC, and NPC, respectively). GPCs, as defined by high *PDGFRA* expression^33^, efficiently generated both immature and mature oligodendrocytes or astrocytes (clusters imOL, maOL, imAstrocytes, maAstrocytes). A smaller population of less defined cells were deemed to be fibroblast-like from high expression of extracellular molecules like alpha-1 type I collagen (*COL1A1*), sharing little expression with remaining clusters.

In vitro, HD hGPCs contained larger NPC clusters than their WT counterparts (**Figure 4D**), and appeared less lineally-restricted than WT, with high *GAD2* suggesting GABAergic interneuronal potential^34^, and only sporadic *OLIG1* expression (**Figure 4E**). Several KZFPs were highly expressed in cells within the integrated NPC cluster, with lower but broader expression across more differentiated glial clusters (**Figures 4F**-4**G**). While overall KZFP expression levels were similar between conditions, the HD GPC and cGPC clusters exhibited a greater proportion of cells expressing primate-specific KZFPs than did WT hGPCs (**Figure S3C**), especially with regards to sporadically expressed KZFPs such as *ZNF492* and *ZNF717*. The mis-timed, heterochronic expression of KZFPs in HD hGPCs thus appears to augur the glial maturational defects associated with mHTT pathology.

### Impaired glial fate commitment in HD hGPCs is highlighted by persistent expression of neural transcription factors and MYST histone acetyltransferase KAT6B

To identify transcriptional abnormalities specific to glial progenitors, we used single-cell RNA-Seq to compare the gene expression profiles of HD and WT GPCs derived from both in vivo and in vitro samples. The most dysregulated genes (|Log_2_FC|>0.15, *p_adj_*<0.05, **Figure 5A**) suggested an HD-associated impairment of oligodendroglial differentiation, suggested by the downregulation of *SOX6, S100B*, and especially *SOX10*, which was suppressed in HD GPCs both in vitro and in vivo, recapitulating earlier findings from bulk expression datasets (**Figure S3D**)^10^. Of note, the downregulation of *SOX10* was associated with the upregulation of *SOX4* and *SOX11*, which can maintain early NPCs in development, but which may also promote neuronal fate in the absence of other SOX factors^35,36^. Accordingly, HD GPCs upregulated several early neural transcripts, including *ARX*, *DLX1,* and *DLX2*, while they exhibited impaired maturation in vivo from downregulation of genes involved in glial maturation, cell–cell and matrix interactions, and synaptic engagement and glutamate uptake, including *S100B, SEMA3E, CNTNAP5, SPARCL1, PSD3, SLC1A2,* and *SLC1A3* (**Figures 5A** and **S3E**)^6,14,37,38^ – all consistent with the impaired glial differentiation of HD hGPCs.

**Figure 5.**
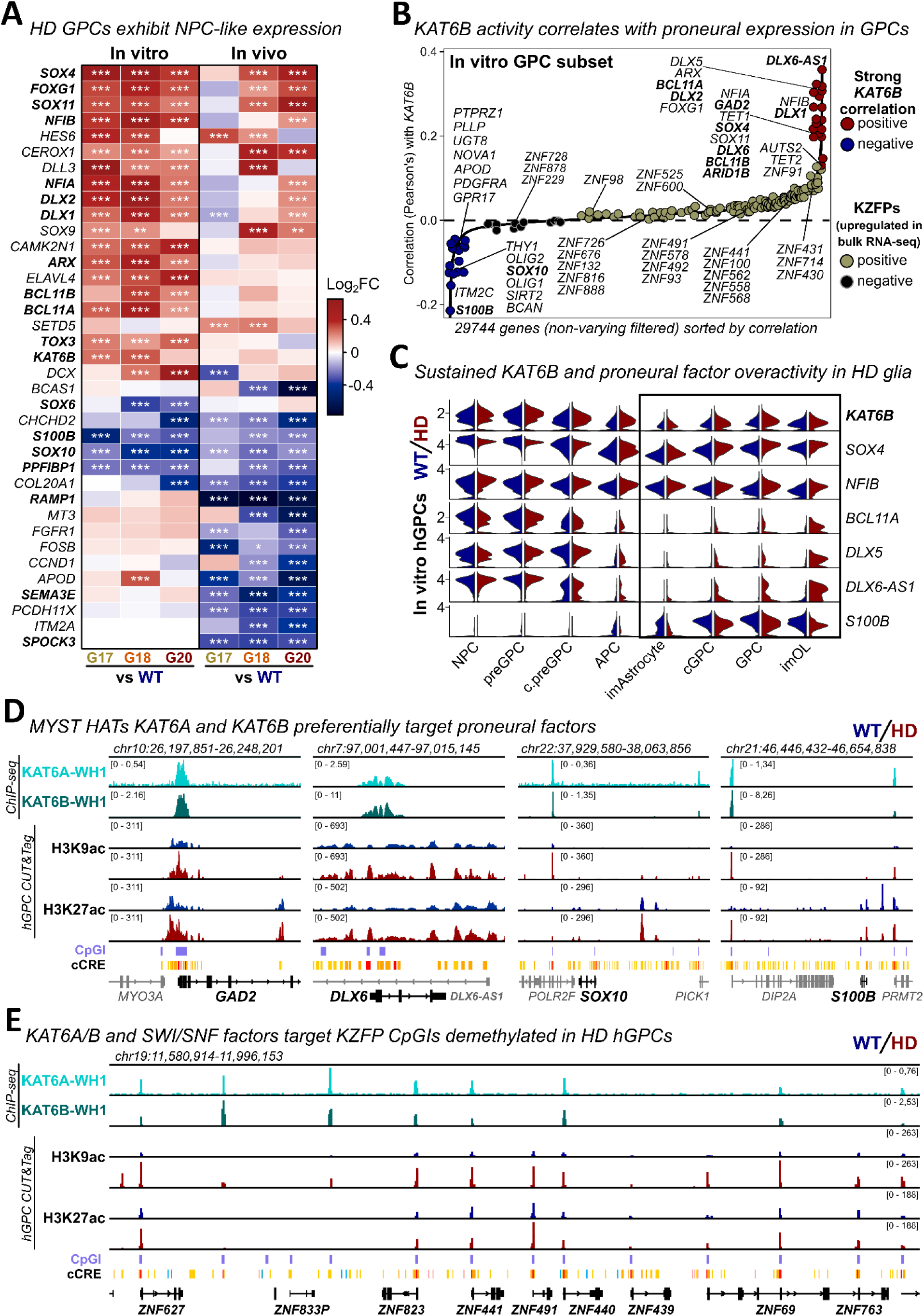
Overactive MYST family histone acetyltransferases KAT6A and KAT6B target KZFPs in HD hGPCs. **A.** Differentially expressed genes identified using *MAST* in the GPC clusters across conditions when comparing each HD line (G17, G18, and G20) to the pooled WT samples. The heatmap lists genes with shared expression patterns deemed to be significant (|Log_2_FC|>0.15 and FDR<0.05). Asterisks indicate significance; *P < 0.05; **P < 0.01; ***P < 0.001. **B.** Pearson’s correlations calculated for each gene within the in vitro GPC subset (WT and HD) show that the expression of glial genes like *SOX10* and *S100B* is anticorrelated to that of *KAT6B*, while neural transcripts like *DLX1, DLX6-AS1,* and *SOX4* are co-expressors. **C**. The expression patterns of MYST family histone acetyltransferase (HAT) *KAT6B* (MORF) and highly correlated neural transcripts are compared between WT and HD clusters identified in hGPCs, and suggest that their overactivity is sustained in HD. **D.** The IGV gene tracks show the binding of MYST HAT KAT6B and its complementary paralog KAT6A for comparison (ChIP-seq datasets from overexpression in HEK293). KAT6A and -B are found at CpGIs of proneural factors, which are highly acetylated at H3 in HD hGPCs, while both are absent from the *SOX10* and *S100B* loci. **E.** KAT6A and KAT6B are recruited to KZFP gene promoter CpGIs on chr19, as shown here at the p13.2 cluster, which exhibit H3K9 hyperacetylation in G20 (HD) hGPCs. The binding data presented is derived from pre-validated and publicly available ChIP-seq assays (as listed in the methods section).

Notably, among the differentially expressed genes associated with impaired glial fate commitment, the MYST-family histone acetyltransferase (HAT) KAT6B (MORF) stood out for its persistent expression in HD hGPCs. In the in vitro GPC subset, *KAT6B* showed the most pronounced and consistent upregulation in HD relative to WT, recapitulating patterns seen in the bulk RNA-seq datasets (**Figures 5A** and **S3D**). To determine whether other epigenetic regulators might similarly contribute to the aberrant chromatin landscape in HD hGPCs, we next examined a focused set of chromatin-modifying enzymes (**Figure S4A**). These included additional HATs; *KAT2A* (GCN5)*, KAT2B* (P/CAF; *p300/CBP-associated factor*), and KAT6A (MOZ), as well as the H3K9 demethylase JMJD1C^39^, the H3K9ac reader CECR2^40^, the H3K27me3 demethylase KDM6B (UTX)^41^, and numerous histone deacetylases (HDACs), which have been of particular interest as targets in HD^42^. Most of these factors showed only modest, inconsistent and largely non-significant changes in HD GPCs relative to WT, reinforcing KAT6B as a main driver of the aberrant acetylation patterns. To place KAT6B within a broader lineage context, we further compared its expression to that of *KAT2A, KAT2B* and *KAT6A* across hGPCs subsets (**Figure S4B**). While *KAT2A* and especially *KAT2B* were preferentially enriched in glial populations (GPC, APC, imAstrocyte, maAstrocyte, imOL, maOL), KAT6A and KAT6B were more prominent in NPCs, consistent with their established roles in neural progenitors^43^. This suggests that it is the selective increase of *KAT6B*, rather than broad-based shifts across HATs or HDACs, which drives the epigenetic and transcriptional glial immaturity of HD hGPCs.

In agreement, the expression of key glial genes found to be downregulated in HD GPCs, including *SOX10* and *S100B*, showed the strongest inverse relationship with *KAT6B* activity in the in vitro GPC subset, as determined by Pearson’s correlation (**Figure 5B**). Conversely, neural TFs, such as *SOX4*, *DLX2, ARX, DLX5*, *DLX6-AS1*, *BCL11B, NFIA,* or *GAD2* were co-expressed with KAT6B and highly upregulated in HD hGPCs (**Figures 5A**-**5C**). Moreover, Homer analysis revealed significant enrichment of DNA binding motifs belonging to DLX and SOX family TFs within hyperacetylated regions (H3K9ac and H3K27ac, **Figure S4C**), underscoring their involvement in over-activation of enhancer-promoter sites in HD hGPCs. Interestingly, the majority of significantly upregulated chr19 KZFPs identified in HD hGPCs (**Figure 3D**) tended to be co-expressed with upregulated *KAT6B*, although these correlations were often modest due to sporadic expression in the GPC subset. Taken together, these data suggest a causal link between the sustained KAT6B expression and early cell-fate determinants in mHTT-expressing hGPCs, alongside delayed and deficient glial differentiation in HD.

### Activated KZFP promoters in HD hGPCs recruit upregulated acetyltransferase KAT6B

To study the recruitment of the upregulated KAT6B, we analyzed publicly available chromatin immunoprecipitation sequencing (ChIP-seq) data for KAT6B and its compensatory paralog KAT6A^43,44^. These data were derived from HEK293 cells overexpressing FLAG-tagged winged helix 1 (WH1) domains in each HAT, which specifically target unmethylated CpGIs, as demonstrated at the *SOX4* promoter (**Figure S4D**)^45,46^. Both KAT6A and KAT6B were highly enriched at promoter-associated CpGIs of additional neural genes – for example, at *DLX6,* and *GAD2*, consistent with the co-upregulation of these transcripts in HD hGPCs. In contrast, they were largely absent from CpGIs at the *SOX10* or *S100B* loci (**Figure 5D**).

We then examined the promoter CpGIs of clustered KZFP genes on chr19, which were both unusually accessible and heterochronically expressed in HD hGPCs, and found that both KAT6A and KAT6B effectively can target these loci (**Figures 5E** and **S4E-S4F**). These findings suggest that sustained dysregulated expression of KAT6B may drive KZFP upregulation in HD hGPCs.

### ZNF98 promotes sustained transcriptional dysregulation in HD hGPCs

To identify upstream drivers of the differential expression signature observed in the HD GPC subset in vitro, we performed motif enrichment analysis (*RcisTarget*) on the significantly upregulated or downregulated genes identified; to do so, we focused on their promoter regions (±10 kb of their TSSs). Among 32 motifs identified from the downregulated gene sets we picked up some of the KZFPs also found to be highly upregulated in HD hGPCs, such as *ZNF208*, *ZNF823*, and *ZNF98* (**Figures 6A** and **3D**). As might be expected due to their repressive function, fewer KZFPs were present among the motifs discovered in the upregulated genes (97 in total, **Table S3**). Interestingly, one of these, ZNF492, is a recent duplication of the primate-specific *ZNF98*^47^ that lacks a functional KRAB domain (**Figure S5A** and **Table S4**), and was suggested to either assist in the activation of the *RPE65* promoter or potentially prevent other repressors from binding^48^. Bulk RNA-seq and CUT&Tag datasets indicated the marked upregulation of both ZNF98 and ZNF492 in HD hGPCs (**Figure S5B**), mirroring the expression patterns of other evolutionary young KZFPs (e.g. *ZNF208, ZNF257, ZNF676,* and *ZNF728* in the 19p12 cluster, **Figure S5C**), although *ZNF98* showed the most consistent upregulation across HD lines (**Figure 3D**). Single-cell RNA-seq revealed that the expanded fraction of NPCs in the HD hGPC cultures were primarily responsible for the over-expression of these KZFPs (**Figures S5D**), thus linking the misexpression of these transcripts to the impaired differentiation of the hGPCs.

**Figure 6.**
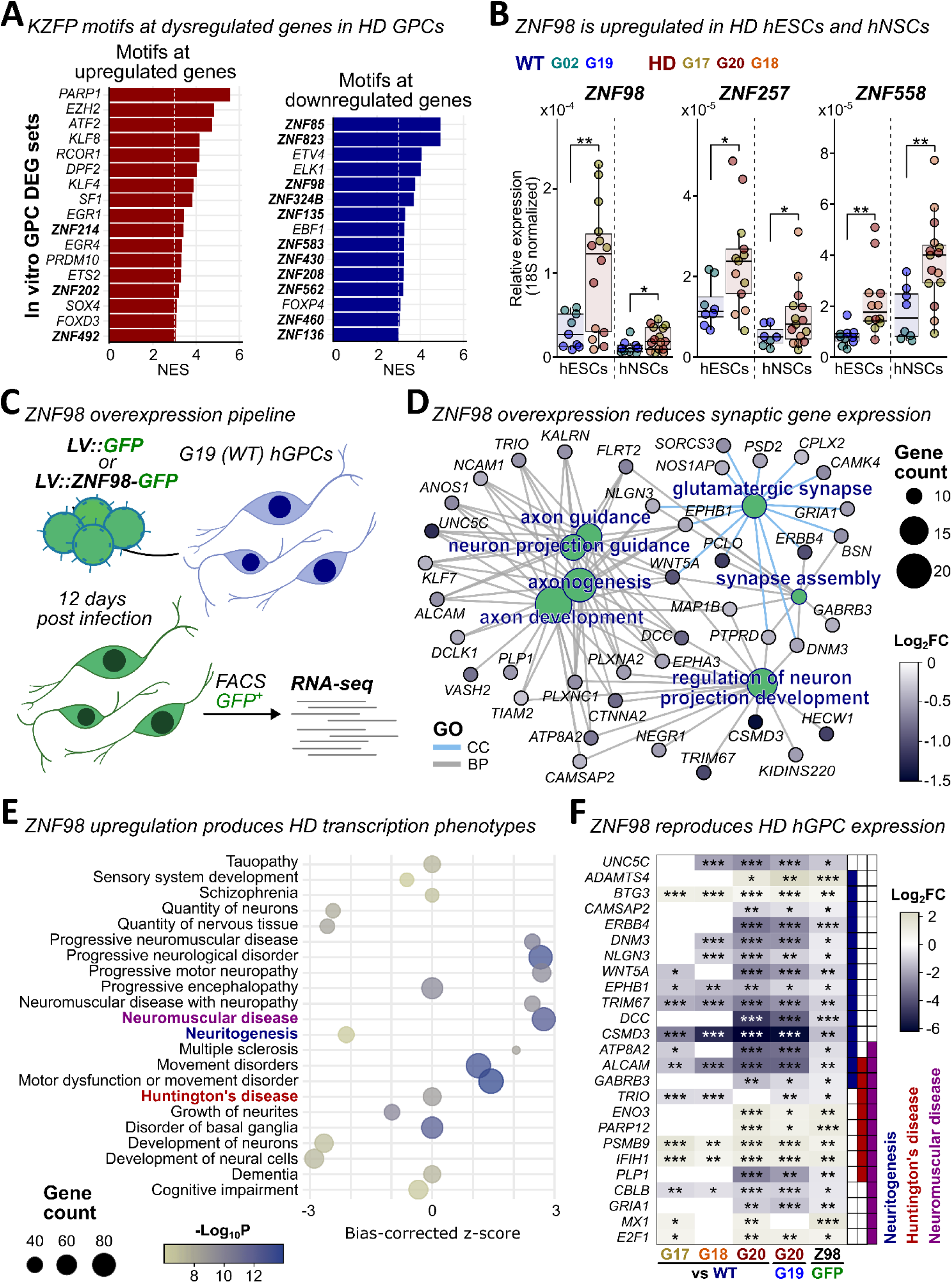
ZNF98 overexpression recapitulates transcriptional signatures of HD hGPCs. **A.** The normalized enrichment scores are shown for significantly enriched transcription factor motifs identified by *RcisTarget* at significantly upregulated (red) or downregulated (blue) genes found in the in vitro HD GPC subset (|Log_2_FC|>0.15 and FDR<0.05). KZFP family factors are shown in bold. **B.** Comparative RT-qPCR analysis of *ZNF98, ZNF257*, and *ZNF558* expression in multiple HD lines (n=3-6/line) at the pluripotent stem cell stage (hESCs) and after 30 days of neural induction (hNSCs) reveals significant differences in KZFP transcript levels between HD and WT groups (Student’s t-test). The scaled y-axes reflect differences in expression magnitude between cell stages. **C.** G19 (WT) hGPCs were transduced with lentiviral constructs expressing ZNF98 and GFP or GFP alone (n=4/group). GFP-positive hGPCs were isolated by fluorescence-activated cell sorting (FACS) 12 days post-infection for RNA-seq analysis. **D.** Gene ontology (GO) enrichment network showing biological process (BP, grey lines) or cellular component (CC, blue lines) terms among significantly downregulated genes following ZNF98 overexpression (*DESeq2*, Log_2_FC<-0.5, p_adj_<0.05). **E.** Ingenuity pathway analysis (IPA) of significantly altered transcripts (*DESeq2*, |Log_2_FC|>0.5, p_adj_<0.05) predicts effects on disease-related and functional categories commonly associated with neurodegenerative disorders, including HD. **F.** Transcriptional changes induced by ZNF98 partially mirror dysregulation observed in HD hGPCs (RNA-seq), particularly among genes contributing to IPA categories such as *Neuritogenesis* (Blue)*, Huntington’s Disease* (Red) and *Neuromuscular Disease* (Purple). Statistical significance is marked by asterisks; *P < 0.05; **P < 0.01; ***P < 0.001.

Since these young KZFPs are reported to be highly active at early stages of development, including embryonic genome activation^22^, we next investigated the relative expression of *ZNF98* in HD embryonic stem cells and neural stem cells (hESCs and hNSCs, **Figure S5E**), compared to their WT counterparts prior to glial differentiation. The upregulated KZFPs *ZNF257* (also from the 19p12 cluster) and *ZNF558* (adjacent to the 19p13.2 cluster) were included for comparison. All three were more highly expressed by HD than WT cells at these earlier stages (**Figure 6B**, **Figure S5D**).

### Expression of ZNF98 in WT hGPCs reproduces aspects of the differentiation block of HD hGPCs

Given the selective expression of *ZNF98* early in glial development, we asked if ZNF98 overactivity in HD hGPCs might trigger the suppression of glial maturation-associated gene expression. To do so, we used RNA-seq to compare the transcriptional profiles of G19 (WT; 172 DIV) hGPCs transduced with lentivirally-expressed ZNF98, to matched cells transduced with a GFP control, 12 days after lentiviral infection (n=4/group, **Figure 6C**, **Figures S6A-C**).

Among 457 differentially expressed genes (p_adj_<0.05, |Log_2_FC|>0.5), gene enrichment analysis revealed the significant downregulation of gene ontologies involved in arborization, synaptic plasticity, and glutamate signaling (**Figure 6D**). The downregulated transcripts included surface receptors critical for contact-mediated signaling and neuron–glia interactions, such as adhesion molecules and netrin signaling factors; *ALCAM*^49^*, EPHB1*^50^*, NCAM1*^51^*, DCC, UNC5C*^52^*, CTNNA2*, *WNT5A*^53,54^ and *ERBB4*^55,56^. Conversely, ZNF98 overexpression led to the upregulation of genes associated with structural plasticity (e.g. *GSN, CD44, VIM*^57-59^) and somatic stem cell properties (*ID3, GSX2, PROM1*^60-62^) (**Figure S6D**).

Ingenuity Pathway Analysis (IPA) of all differentially expressed genes following ZNF98 overexpression predicted decreased activation of neuritogenesis, an ontology that includes genes associated with glial fiber elaboration (**Figure 6E**), further suggesting an impairment in contact-mediated maturation. In addition, disease terms relevant to HD such as progressive neurological disorder and movement disorder were also significantly activated, indicating that ZNF98 overexpression can recapitulate key aspects of HD-associated transcriptional pathology. Indeed, genes dysregulated as a result of ZNF98 overexpression largely mirrored the misexpression observed in HD hGPCs by RNA-seq (**Figure 6F**), indicating that aberrant KZFP expression was sufficient to reproduce much of the glial transcriptional dysregulation of HD.

## DISCUSSION

Glial differentiation failure in Huntington disease may comprise a significant contributor to both its clinical phenotype and disease progression. Human HD glial progenitor cells (hGPCs) show cell-intrinsic defects characterized by delayed and deficient astrocytic and oligodendrocytic differentiation, as reflected in the suppressed expression of key regulators of glial lineage, such as SOX10, MYRF, and TCF7L2^10,15,17^. Accordingly, both human and animal studies reveal early white matter abnormalities and glial maturation defects in HD, which likely serve to exacerbate neuronal circuit dysfunction^4,12,14,63^. Yet despite these provocative indications that glial differentiation failure may be a central participant in HD pathogenesis, its mechanistic basis has remained largely underexplored. In this study, we examined the epigenetic landscape of HD glial progenitor cells and discovered that their chromatin was differentially accessible at the promoters of KZFP genes clustered on chr19, leading to the persistent expression of these KRAB-domain transcriptional repressors. The HD hGPCs exhibited delayed and deficient glial differentiation, exemplified by their suppression of *SOX10* and *S100B,* with attendant deficits in both astrocytic and oligodendrocytic maturation^10,17^. The upregulation of KZFP gene expression was strongly associated with H3K9 hyperacetylation, which coincided with elevated levels of the MYST family histone acetyltransferase KAT6B (MORF); the resultant hyperacetylated chromatin state was associated with the maintenance of HD hGPCs at an immature stage, at the expense of terminal glial differentiation. Accordingly, we found that the forced overexpression in WT hGPCs of ZNF98, a primate-specific KZFP, was sufficient to recapitulate key aspects of HD hGPC transcriptional dysregulation.

Glial differentiation requires extensive chromatin remodeling to silence neural and neurogenic transcription programs, and is typically associated with a global reduction in histone acetylation and heterochromatinization – a process conserved across phylogeny^64,65^. The hyperacetylation of HD hGPCs may therefore disrupt their glial maturation, which depends on histone deacetylase (HDAC) activity to suppress WNT/β-catenin signaling antecedent to glial fate commitment^66-68^. Accordingly, HDAC inhibition can impair the maturation of both oligodendrocytes and astrocytes, so much so as to promote neuronal gene expression in glioma cells^69-71^. Additional impediments to glial maturation in HD hGPCs include their reduced expression of TCF7L2 previously documented^10^, which otherwise acts to drive oligodendrocyte differentiation and myelinogenesis by counteracting WNT signaling^16^. Indeed, the persistent WNT signaling enabled by histone hyperacetylation in HD hGPCs might be expected to maintain NPC identity at the expense of oligodendroglial and astroglial development^72-75^, just as we observed.

In line with the H3K9 hyperacetylation of HD hGPCs in vitro, we found that these cells significantly upregulated expression of KAT6B. Although our study did not directly assess non-histone acetylation targets, prior research has shown that MYST HATs (e.g., MOZ/KAT6A, KAT5, KAT7) enhance WNT signaling by stabilizing β-catenin and activating TCF-mediated transcription. HD hGPCs may therefore increase their WNT/β-catenin signaling via both chromatin relaxation and DNA demethylation at several WNT pathway promoters (**Figure S2D**), as well as by the activation of β-catenin^76,77^. In this regard, TFs including DLX2, ARX, SOX4, and SOX11 – which act as both promoters of WNT/β-catenin signaling^78-80^ and suppressors of oligodendrocyte development^81,82^ – were upregulated in HD GPCs in vitro, paralleling the increased levels of KAT6B/MORF. These observations are consistent with the roles of KAT6A/MOZ and KAT6B/MORF in activating the DLX and SOX gene families, and in promoting the maintenance of NPCs^83-85^. As such, these HATs may prove promising targets for counteracting the histone hyperacetylation-linked failure of both astrocytic and oligodendroglial differentiation in HD^72-75^.

Of note, although HATs possess intrinsic DNA-binding domains, their genomic targets in the context of glial development remain poorly characterized. Like HDACs, HATs are recruited by TFs such as MYC, which directs KAT2A to activate enhancers^86^. Notably, in our maturing glial populations, KAT2A and KAT2B (PCAF) displayed selective activities, diverging from the expression patterns of KAT6A and KAT6B (**Figure S4B**), suggesting these HATs, by recruiting chromatin remodeling complexes, can facilitate the opening of heterochromatic regions and influence developmental plasticity. Consistent with this, HD hGPCs upregulated several DNA-binding factors known to recruit such complexes to enhance chromatin accessibility and regulate cellular reprogramming, including SOX4, PBX1, DLX1, and BCL11B^81,87-89^. Interestingly, we noted that BCL11B binding sites intersecting not only with KAT6B and the aberrantly accessible and hyperacetylated KZFP promoters on chromosome 19, but several core components of the neural-progenitor BAF (npBAF) SWI/SNF complex – namely BRG1 and ARID1B (**Figures S4E-S4F**), which are essential for maintaining NPC self-renewal^90^. Both hyperacetylation and increased expression of EVF2 (*DLX6-AS1,* **Figure 5D**) are known to stabilize SWI/SNF binding, but impair consecutive rounds of ATP-dependent chromatin remodeling^91-94^, whereas ARID1B-haploinsufficient stem cells fail to exit early pluripotency^95^. This suggests that the npBAF complex may be overly retained in HD hGPCs and prevent efficient progression from NPC-like states, but whether this is also the case in HD hGPCs remains to be experimentally validated.

The aberrant activation of KAT6B and its associated pioneer factors in HD hGPCs appears to further enhance chromatin accessibility and KZFP gene activation, recapitulating molecular features of early developmental and reprogramming states. Such widespread activation of primate-specific KZFPs mirrors patterns observed during embryonic genome activation, or following somatic cell reprogramming to stem cell pluripotency^22,96^. This is exemplified by the premature induction of KZFPs such as the primate-specific ZNF98, which upon overexpression in healthy hGPCs led to upregulation of *GSX2* and *ID3,* as well as of the early neural transcripts *PROM1*, *VIM*, and *CD44*. This indicates that ZNF98 overactivity can also promote a shift towards less mature, stem-like phenotypes in HD hGPCs^57-59,62,97^. Complementary findings have been made following the antagonism of other primate-specific KZFPs in NPCs; ZNF675 and ZNF558, which resulted in premature neuronal maturation^98,99^. Notably, increased expression of *PROM1, VIM,* and *CD44* have all been observed in human HD astrocytes^14,63,100^, while a recent multiomic study of HD patient-derived astrocytes linked the dysregulation of CD44 to *mHTT* CAG repeat length^101^. However, while increased *CD44* expression may highlight the acquisition of progenitor features, its upregulation may also reflect increased inflammatory signaling^102^, as suggested by a number of co-expressed interferon-associated factors (**Figure S6D**).

In contrast to the upregulation of early neural genes in hGPCs in response to ZNF98, the expression of a broad set of genes involved in cell-cell interactions, axon guidance, and synapse assembly fell, mirroring aspects of the expression signature of HD hGPCs. Transcripts down-regulated following ZNF98 expression included several netrin pathway signaling components such as *UNC5C, UNC5D* and *DCC*, whose suppression suggested that ZNF98 might play a critical role in gliovascular as well as gliosynaptic interactions^103,104^. Dysfunctional netrin-mediated oligodendrocytic contact might then be expected to impair axonal adhesion and thereby compromise myelin integrity^105^. In addition, netrin pathway dysregulation may contribute to the abnormal morphologies observed in both HD hGPCs and astrocytes^10,12,14^, given netrin’s role in process branching and self-avoidance^106,107^. Furthermore, the reduced expression of other mediators of cell-cell interaction, including the UNC5/latrophilin ligand *FLRT2, as well as NLGN3* and *DNM3* (**Figures 6D** and **6F**), link ZNF98 to the impaired synaptic engagement, plasticity, and signaling of HD hGPCs^108-110^. Of note, both HD and schizophrenia-derived hGPCs downregulate *NLGN3, DNM3* and the netrin ligand *LRRC4C* (**Figure S1E**), as well as both *NTNG1* and latrophilin 3 (*ADGRL3*)^111^. This observation suggests that the reduced expression of these mediators of glial-glial and glial-neuronal interaction may comprise a feature of glial pathology common to these disorders. As such, ZNF98-regulated transcription thus predicts disruption of glial contact-mediated signaling and phenotypic maturation.

Of note, our findings in human HD glia contrast with several studies in rodent cellular models of HD, which have reported that mHTT inhibits HATs such as CBP/CREBBP and P/CAF/KAT2A^112-114^; these data have prompted the assessment of histone deacetylase (HDAC) inhibitors as a therapeutic approach^115-117^. Yet while such HDAC inhibition yielded functional benefit in rodent HD models, clinical trials of HDAC inhibitors in HD patients failed to demonstrate efficacy^42,118,119^. Importantly, our human HD glial cells display a notably different epigenetic profile than their rodent counterparts, with only minor upregulation of HDACs (**Figure S4A**), suggesting a species-linked difference in epigenetic regulation that may explain the disconnect between preclinical and clinical outcomes to HDAC inhibition. In this regard, Narayan et al.^120^ recently reported the hyperacetylation of histones H3 and H4 (including H3K9ac) in brain cells of both HD patients and a transgenic HD sheep, complementing the hyperacetylation seen here in HD hGPCs. Together, these data suggest some limitations to the use of rodent models of HD in predicting the therapeutic effects of epigenetic modifiers. Yet beyond these species-linked differences in epigenetic dysregulation, a systematic review of re-analyzed RNA-seq datasets from human HD samples, including both patient tissues and stem cell-derived cells^121^, identified a number of mis-expressed epigenome remodelers, but with variability across cell type. Such variability highlights the context-dependent nature of epigenetic regulation, and hence the critical influence of cellular composition and brain region on HD signatures. As such, rodent and bulk tissue studies may oversimplify or miss key epigenomic alterations in human HD, especially those referable to specific cellular phenotypes. Together, these findings suggest the need for human-centric, single-cell multi-omic analyses of defined cellular subpopulations when assessing the transcriptional architecture – and therapeutic vulnerabilities of HD^122^.

Together, these data reveal that the upregulation of MYST family HAT KAT6B in HD hGPCs promotes profound H3K9 hyperacetylation in HD hGPCs, at recently evolved KZFP clusters found across chr19, leading to the aberrant expression of a set of KRAB domain finger repressor proteins. These transcriptional and epigenetic abnormalities are linked closely to the abnormally prolonged maintenance of progenitor state, at the expense of glial development and function. This effect was exemplified by the overexpression of the primate-specific ZNF98 in wild-type hGPCs, which reproduced much of the expression signature of HD-derived hGPCs.

## METHODS

### Human glial progenitor cells

Bipotential human glial progenitor cells (hGPCs) were prepared as previously described from human embryonic stem cells (hESCs)^123^. Briefly, the hESCs were passaged onto human laminin-521 (Corning, CB40222) using gentle cell dissociation reagent, and maintained in mTeSR (Stemcell technologies, 100-0485 & 100-0276) in a 5% CO2 incubator at 37 °C. The hESCs were not passaged more than 7 times; following each passage, cells were lifted for embryoid body (EB) formation and hGPCs were prepared as described^123^. All hGPCs were assayed after 160-200 days in vitro (DIV); cells were routinely checked for contamination and mycoplasma. Three lines each of WT and mutant HTT-expressing (HD) hESCs were used, derived from either healthy wild-type (WT: GENEA02 and -19; also WA09) or Huntington’s disease (HD: GENEA17, -18, and -20) blastocysts. GENEA19 (G19; WT, 18Q) and GENEA20 (G20; HD, 48Q) are female fraternal siblings. The number of CAG repeats in *HTT* on each allele is as follows:

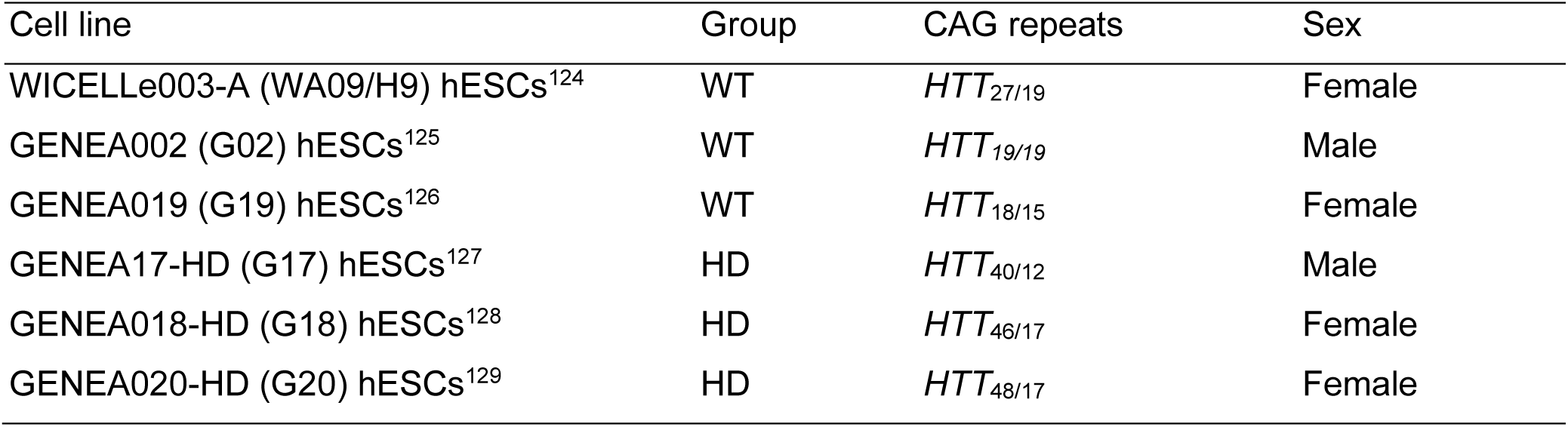

### Animals

Homozygous *rag2* null immunodeficient C3H mice (Taconic, 000602-M) were crossed with the myelin-deficient (*MBP^shi/shi^*) *shiverer* mice (The Jackson Laboratory, 001428) to generate *shiverer x rag2^-/-^* (SR2) mice as described^130^. The mice were maintained on a 12:12 hour light cycle in pathogen-free housing, with a controlled temperature and humidity, and were fed *ad libitum*. The experiments have been approved by the University of Rochester’s Committee on Animal Resources (UCAR).

### Cell transplantation and engraftment

hGPCs were prepared for transplant at DIV160 as described^130^ with collection and gentle dissociation in Ca^2+^/Mg^2+^-free HBSS^-/-^ (Thermo Fisher Scientific, 14170112). The HBSS/cell suspensions (10^5^ cells/µL) were injected bilaterally into 2 sites/hemisphere (5·10^5^ per site) in the anterior and posterior corpus callosum (CC) in neonatal (P1-P3) SR2 mice. Each litter was weaned between days 21-28; mice were euthanized by CO_2_-inhalation 19 weeks post transplant, then processed for scRNA-seq.

### Immunocytochemistry

Cultured cells were fixed in 4% paraformaldehyde and used for immunocytochemistry (ICC). Fixed cells were then stained in two consecutive rounds using primary and secondary antibodies in a permeabilization buffer with a blocking agent; buffered saline (DPBS) with 1% bovine serum albumin (BSA) and 0.3 % Triton-X. The following primary antibodies were used; anti-OCT4 (R&D Systems, MAB17591, 1:100) and anti-NANOG (Abcam, ab21624, 1:100) in hESCs; anti-SOX1 (R&D Systems, AF3369, 1:100) and anti-PAX6 (Biolegend, 901301, 1:300) in neural stem cells (hNSCs) at DIV30; anti-Olig2 (Merck Millipore, MABN50, 1:1000), anti-hPDGFRA (Cell signaling, 5241, 1:300) and anti-GFAP (Invitrogen, 13-0300, 1:500) in hGPCs (DIV160-180). Secondary antibodies were used at dilution 1:500 and are listed with all antibodies in the resource table, while DAPI was used for nuclear staining (Sigma Aldrich, D8417, 1µg/mL).

### Flow cytometry and cell sorting

Surface markers of hGPCs were assessed by flow cytometry. Adherent cells were lifted using 50% Accutase (Stemcell Technologies, 07922) in PBS and triturated gently for single cell suspension in DMEM with 100 U/mL DNase I (Sigma Aldrich, D4263), then washed in DMEM/F12. Cells were resuspended in ice-cold wash buffer (PBS pH7.2 with 0.5% BSA and 2mM EDTA) at approx. 10^6^ cells/mL and stained for 20 minutes on ice in the dark with slight agitation. The following conjugated antibodies were used: CD140a-PE (BD Bioscience, 556002, 1:10), CD44-APC (Miltenyi Biotec, 130-113-331, 1:200) and A2B5-APC (Miltenyi Biotec, 130-093-582, 1:50). All remaining steps were carried out on ice. The stained cells were rinsed once in wash buffer, resuspended in DMEM/F12 with 20U/mL DNase I, and passed through a 35µm cell strainer, before adding DAPI (Sigma Aldrich, D8417, 1µg/mL) used to assess cell viability. Cell fluorophore intensities in the final suspensions were measured in a FACSAria IIIU (Becton Dickinson) and analyzed with the FACSDiva (BD Biosciences) software. For Fluorescence-activated cell sorting (FACS) of DAPI^-^ and CD140^+^ hGPCs or GFP^+^ hGPCs (following infection with LV-ZNF98-FLAG-GFP), positive populations were sorted into ice-cold DMEM in falcon tubes, which had been pre-coated for a minimum of 15 min with coating buffer (PBS pH7.2 with 3.375 % BSA) at RT.

### DNA methylation array

Four sets of hGPCs induced from G19 (WT) or G20 (HD) hESCs were CD140-sorted at DIV180 for genomic DNA extraction. The DNA was bisulfite-treated and hybridized on a Infinium HumanMethylation 450 BeadChip array (Illumina, WG-314). Following sequencing, the Illumina microarray expression data (.idat) files were normalized by the stratified quantile normalization procedure preprocessQuantile from *minfi*^131^. 439,972 probes were retained following filtering of probes exhibiting single nucleotide polymorphism, cross-reactivity or low confidence probes (pval>0.01). Differential methylation analysis was performed on the matrix of log2 ratio of the intensities of methylated versus unmethylated probes (M-values). Briefly, a linear fitted model was constructed with *∼0+condition* design, which was then processed through a contrast fit (HD-WT). The linear model was subsequently inputted into an empirical Bayes moderated t-statistics test, the eBayes function from *limma*^132^, to look for probes with evidence of differential methylation. A pre-compiled hg19 database from Bioconductor was used for annotation (IlluminaHumanMethylation450kanno.ilmn12.hg19) and lifted to hg38. *ChIPseeker*^133^ was used for further gene and feature annotation of DM sites.

### Assay for transposase-accessible chromatin sequencing (ATAC-seq)

ATAC-seq libraries were constructed as described previously^30^. In brief, 50k nuclei were isolated from CD140-sorted G19 (WT) or G20 (HD) hGPCs (n=2) at DIV180 using cold lysis buffer. Immediately after, tagmentation was carried out, followed by Nextera DNA library preparation (Illumina, 15028212 and 20034197). Library fragments were amplified with NEBnext PCR reagents (New England Biolabs, M0541) and purified with a MinElute kit (Qiagen 28004), before being sequenced (Illumina, HS2500). Low quality reads were filtered out with *fastp* and then aligned to the human genome (hg38 assembly) with *bowtie2*^134,135^. Duplicate and low-quality reads (MAPQ<30), or reads aligning to blacklisted regions were filtered out using *samtools* and Picard^136,137^, before fragment enrichment was assessed with *Genrich* (https://github.com/jsh58/Genrich) using the default parameters. Following peak calling, consensus peaks were defined using *soGGi*, resulting in 31,232 unique WT regions and 18,531 HD regions, and 49,615 shared regions^138^. Finally, all 99,378 consensus regions were evaluated for differential accessibility with *DESeq2* and annotated with *ChIPseeker^133,139^*.

### Cleavage under targets and tagmentation (CUT&Tag) sequencing

Nuclei were isolated from approx. 300k hGPCs at DIV160-180 using a gentle approach described previously^23^, and subjected to CUT&Tag^140^, using the reagents and protocol (v1.5) from Epicypher. The following primary rabbit antibodies were used (0.5µg/reaction); IgG, Tri-Methyl-Histone 3 (Lys4) and Acetyl-Histone 3 (Lys9 and Lys27) from Cell Signaling (2729S, 9751S, 9649S, 8173S, respectively) and Tri-Methyl-Histone 3 (Lys9) from Active Motif (39065). CUT&Tag libraries were submitted for 2x75nt paired-end sequencing on the NextSeq 550 system (Illumina) at the University of Rochester Genomics Research Center (GRC). Raw sequencing reads were filtered with *fastp^134^*and library quality was further assessed using fastqc^141^. Libraries were then aligned to the human genome (assembly hg38) using bowtie2^135^ with the parameter: --very-sensitive-local --no-unal --no-discordant --no-mixed --dovetail -N 1 -X 1000. Unmapped reads and low quality reads (MAPQ < 12) were filtered out with *samtools* while duplicates were flagged and removed using *Picard^136,137^*. From filtered .bam files, .bedgraph and .bigwig files were generated with *deeptools* bamCoverage using RPGC normalization with the genome coverage calculated for each histone mark type^142,143^. For peak visualization in integrative genomic viewer (*IGV*) software or *deeptools* heatmaps, the median .bigwig signals were calculated from sample replicate .bedgraphs using *bedtools* and *UCSC* tools^144^. Broad peak calling was performed with *MACS2* and used for differential peak enrichment analyses WT and HD samples in *DiffBind*, while *ChIPseeker* was used for peak annotation^133,145,146^.

### Single-cell expression captures

Cells were captured for single-cell RNA sequencing (scRNA-seq) originating either from cell suspensions prepared directly from lifted hGPCs in vitro or tissue-derived cells from transplanted 19 weeks old SR2 mice. Following microdissection of the chimeric corpus callosum, samples were stored in cold HBSS^-/-^before being dissociated in 0.5mL PIPES solution (120mM NaCl, 5mM KCl, 25mM glucose and 20mM PIPES) and 0.5mL with 10units of filtered Papain (Worthington, LS003126. Activated with EDTA/cysteine-HCL as described by the manufacturer) and 60 Kunitz units of DNase I (Sigma Aldrich D4263) a 37°C in an oscillator for 50min with gentle trituration every 20min. The dissociation reaction was stopped with 1.2mL ovomucoid protease inhibitor with BSA (Worthington, LK003182), gently triturated with a P1000 pipette until a single cell slurry was obtained. The cells were washed and centrifuged in MEM (ThermoFisher, 11095080) for 10min before being resuspended and filtered (35µm mesh) directly into FACS tubes (Fisher Scientific, 08-771-23) for sorting and DAPI was applied (1µg/mL) for viability. The DAPI-negative populations (80-95%) were sorted into 0.5mL LoBind Eppendorf tubes with 20µL DMEM. For the captures (n=2-4/category), the Chromium next GEM single-cell 3’ reagents kit v3.1 was used (10X genomics, PN-1000121) following the manufacturers user guide (rev. D). Single cell suspensions were adjusted to 4·10^5^ cells/mL and about 16·10^3^ cells were loaded for each well of chip G for a target recovery of about 10·10^3^ cells per capture. Following reverse transcription and GEM cleanup, library preparation was carried out by the University of Rochester Genomics Research Center (GRC) and sequenced on a NovaSeq 6000 system (Illumina).

### Single-cell data processing and differential expression analyses

Sequence reads were aligned using STARsolo^147^ to a custom reference constructed by concatenating human (GRCh38 Ensembl 106) and mouse (GRCm39 Ensembl 106) genomes and filtering for transcript types: “protein_coding”, “lncRNA”, or “miRNA”. Alignment was carried out with the following flags: “--twopassMode Basic --outSAMtype BAM Unsorted --readFilesCommand zcat --soloType CB_UMI_Simple --soloCBwhitelist 3M-february-2018.txt --soloUMIlen 12 --soloUMIfiltering MultiGeneUMI --soloCBmatchWLtype 1MM_multi_pseudocounts --limitSjdbInsertNsj 2000000”. Following alignment, the STARsolo-filtered UMI count matrix was split into components containing either mouse or human genes. For each species the respective count matrix was filtered for cell barcodes that possessed at least 1000 unique genes expressed and less than 10% mitochondrial gene expression for further analysis. Raw UMIs from cells passing the cut-off criteria in all samples were merged into a matrix and used as input for data integration. Data were integrated while accounting for line, capture, percent mitochondrial content, number of unique features, and number of UMIs as covariates with *scVI^148^*. Cellular architecture and heterogeneity were visualized via UMAP^149^, and leiden cell clusters determined from the integrated model normalized values using *scanPY*^150^. For differential expression analysis, gene sets were first filtered for appreciably expressed genes (10% detection in any comparison group). Differentially expressed genes were identified using MAST, with cell line and scaled number of unique features as fixed effects^151,152^. Multiple comparisons were controlled using the false discovery rate method. Genes with an FDR < 0.05 and an absolute Log_2_-fold change > 0.15 were deemed significant. All genes significantly upregulated or downregulated in at least two disease lines (G20 and G18, G20 and G17, or G18 and G17) compared to pooled the WT GPC subsets were submitted to *RcisTarget* to scan the ±10kb region surrounding their TSSs for motif enrichment, using the default normalized enrichment score (NES) threshold of 3 and motif v10 annotations and rankings databases^153^.

### RT-qPCR analysis of KZFP expression in stem and neural stem cells

RNA was extracted directly from human embryonic stem cell (hESC) cultures, or from neural stem cells (hNSCs) at DIV30 of our glial induction protocol with the RNeasy mini extraction kit (Qiagen, 74104). Per quantitative PCR reaction, 10ng of RNA was converted into cDNA using a TaqMan kit with MultiScribe reverse transcriptase (ThermoFisher, N8080234) and prepared with PowerUp SYBR green master mix (ThermoFisher, A25741). Reactions were set up using the primer pairs listed in the resource table.

### Lentiviral expression of ZNF98

The whole endogenous coding sequence of *ZNF98* including a 3’-terminal FLAG tag was synthesized (Bio Basic, gene synthesis) and lifted into the pTANK-CBh-GFP-WPRE (control) lentiviral vector to generate pTANK-CBh-ZNF98-FLAG-GFP-WPRE. The *CBh* promoter is a strong CMV enhancer/chicken β-actin hybrid (CBh) promoter. The plasmid further contains a T2A self-cleaving peptide between ZNF98 and GFP. Lentivirus was prepared based on the protocol from Didier Trono’s lab using the 2^nd^ generation envelope (pLP-VSV-G/pMD2.G) and packaging (psPAX2) plasmids (Addgene, #12259 and #12260, respectively). The functionality of lentiviral particles was confirmed by a small virus prep; transfection of HEK293FT cells (Thermo Fisher Scientific R70007) in a 35mm well with PEI reagent at 90% confluency (Serochem, PRIME-AQ100) at a ratio 4:1 to 2µg of total plasmid DNA (equimolar amount of each plasmid) in 200µL Opti-MEM (Thermo Fisher Scientific 31985062), which was added to the cells dropwise; replacement of transfection medium after 18hrs; transfer of 1mL of virus medium following another 24hrs to another 35mm well with HEK293FT; fixation, permeabilization and staining of cells 10 days post infection as described above using primary antibodies against GFP (Thermo Fisher Scientific, A10262) and FLAG (Sigma-Aldrich, F1804). Concentrated lentivirus was prepared in an upscaled version of the small preparation described above; HEK293FT cells were transfected in six 150mm culture dishes at a ratio 4:1 to 60µg of total plasmid DNA in 6mL Opti-MEM; replacement of transfection medium after 18h; collection of lentivirus supernatant on the two consecutive days (24-48hrs, stored on ice in refrigeration), pre-clearance of cell debris (2000rpm/10min), and ultracentrifugation (20000rpm/3hrs) at 4°C. Lentiviral particles were recovered in 350-400µL of Opti-MEM after 15min of gentle agitation (150-200rpm) and stored at -80°C before use. Lentiviral titers were calculated using the qPCR Lentivirus Titer Kit from Applied Biological Materials (ABM, LV900). G19 hGPCs at DIV160 were transduced for a calculated MOI of 1 and fresh growth medium was added following 18hrs of incubation. The hGPCs were maintained for 12 days before, GFP-positive hGPCs were sorted for RNA-seq.

### RNA-seq in ZNF98-overexpressing hGPCs

RNA was extracted from GFP-sorted G19 (WT) hGPCs infected to express the ZNF98-FLAG-T2A-GFP, or GFP alone (n=4/group) using the RNeasy micro extraction kit (Qiagen, 74004). The RNA samples were submitted to University of Rochester Genomics Research Center (GRC) for quality assessment, NGS library preparation, and sequencing on a NovaSeq 6000 system (Illumina).

### RNA-seq data processing and differential expression analysis

Raw FASTQ files were trimmed and adapters removed using *fastp* and aligned to GRCh38 using Ensembl 106 gene annotations via *STAR* in 2-pass mode across all samples, and quantified with *RSEM* version 1.3.1^134,147,154^. Subsequent differential expression analysis was carried out using *DESeq2^139^*. These data were then further filtered by meaningful abundance, defined as a median TPM (calculated via RSEM) of 1 in at least 1 group for downstream analyses. Genes with adjusted p-values < 0.05 and Log_2_-fold changes 0.5 were deemed significant and were submitted to ingenuity pathway analysis (IPA, Qiagen). Top enriched categories among canonical diseases restricted to neurological disorders were retained for visualization. Previously generated RNA-seq data from our group using bulk RNA was also used for our initial investigations of KZFP expression in WT and HD hGPCs^10^.

### Chromatin immunoprecipitation sequencing (ChIP-seq) datasets

The following pre-validated ChIP-seq binding datasets were gathered from ChIP-atlas^44^ (GEO/DDBJ study and sample references are included).

- KAT6A-WH1-FLAG rep1; HEK293T (transient transfection); DRA014290/DRX366108
- KAT6A-WH1-FLAG rep2; HEK293T (transient transfection); DRA014290/DRX366109
- KAT6A-WH2-DPF-FLAG rep1; HEK293T (transient transfection); DRA014290/DRX366110
- KAT6A-DPF-FLAG rep1; HEK293T (transient transfection); DRA014290/DRX366111
- Input control; HEK293T; DRA014290/DRX366112
- Empty FLAG vector rep1; HEK293T (transient transfection); DRA015383/DRX411441
- KAT6B-WH1-FLAG rep1; HEK293T (transient transfection); DRA015383/DRX411443
- KAT6B-WH1-FLAG rep1; HEK293T (transient transfection); DRA015383/DRX411444
- BCL11B rep2; primary TALL-1; GSE165209/GSM5024541
- RUNX1; HEK293; GSE85524/GSM2467011
- ARID1B, K562, GSE92083/GSM2424018
- SMARCA4 rep1, human NPCs, GSE122631/GSM3494631

## QUANTIFICATION AND STATISTICAL ANALYSIS

Unless otherwise stated, data were analyzed and visualized using R in RStudio 2024.09.1+394 "Cranberry Hibiscus" Release. Unpaired t-tests were used to compare two groups. Quantitative results are shown as mean ± S.D., with statistical significance accepted at p<0.05.

## RESOURCE AVAILABILITY

### Lead contact

Further information and requests for data should be directed to and will be fulfilled by the lead contact, Steve Goldman.

### Materials availability

Any requests for materials and reagents should similarly be directed to the lead contact, Dr. Goldman.

### Data and code availability

- All in vivo histological data in this paper will be made available upon request by the lead contact
- New code was generated in this study, the scripts for which are available on Github at: https://github.com/CTNGoldmanLab/HD_GPC_KZFP
- The sequencing datasets reported in this manuscript have been deposited to GEO, accession number GSE306143.
- Any further information required to reanalyze any data reported in this paper are also available from the lead contact.

## Author Contributions

PB and SAG designed the study; PB performed vector cloning and lentiviral preparation, immunocytochemistry, flow cytometry, CUT&Tag, and RT-qPCRs. PB and RF raised all hGPCs; SJS performed the cell transplantations and dissections; DS and BM retrieved hGPCs from dissected tissue and performed scRNA-seq captures; JNM and PB analyzed the scRNA-Seq data; RF and NPTH prepared and analyzed the ATAC-seq and DNA methylation data; PB and SAG analyzed all data and wrote the manuscript, to which all authors contributed.

## Acknowledgements

Supported by the Lundbeck Foundation, the Novo Nordisk Foundation, NIH/NIA, the Adelson Medical Research Foundation and the Olav Thon Foundation. We thank the Genomics Research Center of the University of Rochester Medical Center for support.

## SUPPLEMENTAL INFORMATION

**Figures S1-S6,** and **Tables S1** and **S4**.

**Table S2.** Aberrant availability at chr19 KZFP genes in HD hGPCs (Excel table).

**Table S3.** Transcriptional analyses of HD hGPCs (Excel table).

