## Supplemental Figures S1-S6 and Tables S1-S4 for "Histone hyperacetylation-linked upregulation of KRAB zinc finger proteins impedes glial differentiation in Huntington’s disease"

**SUPPLEMENTARY FIGURES**

**Figure S1**

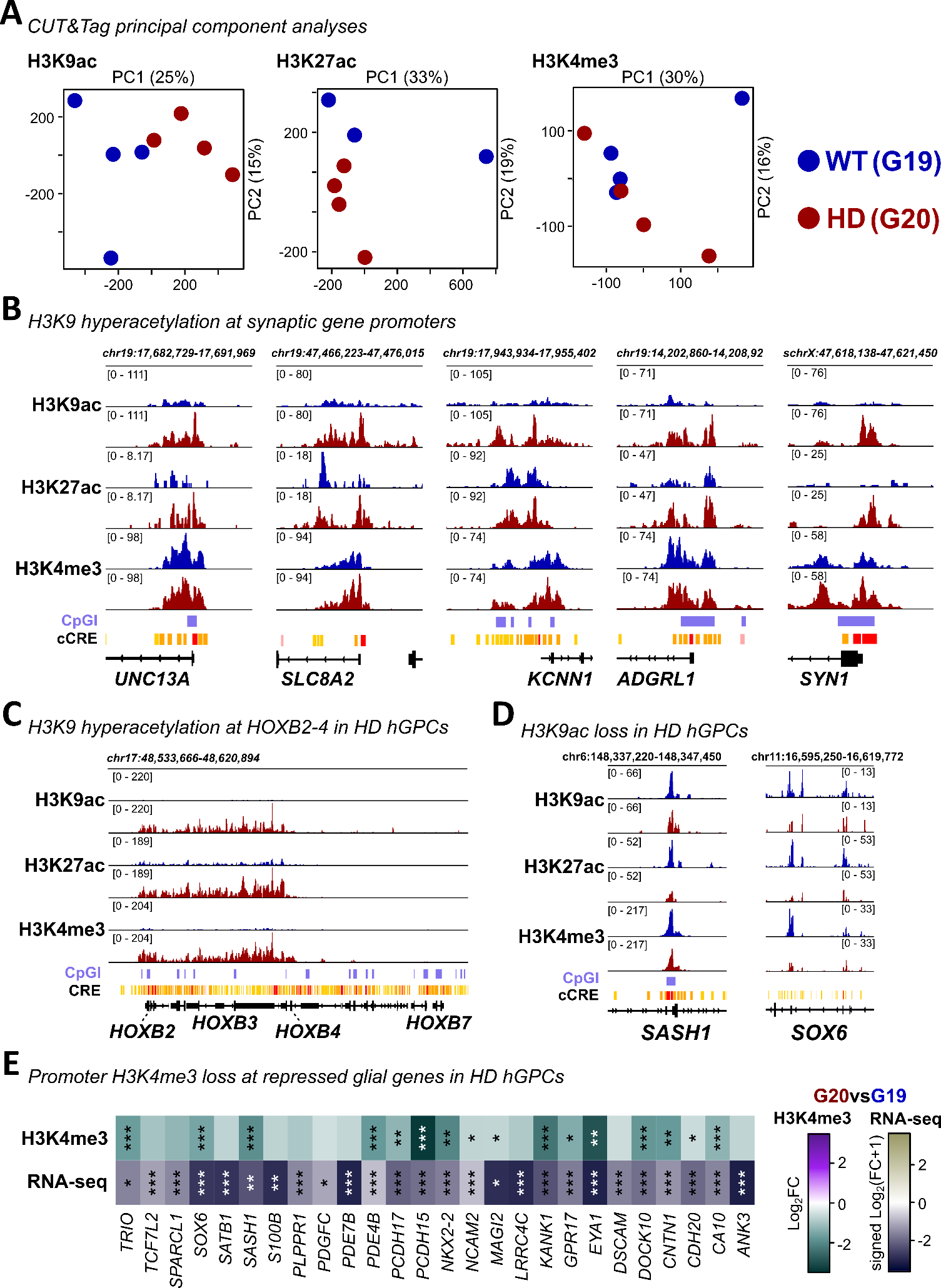

**Figure S1. Chromatin profiling reveals H3K9 hyperacetylation in HD hGPCs, related to Figure 1.**

**A.** Principal Component Analysis (PCA) shows the segregation of histone mark CUT&Tag hGPC samples from sibling lines G19 (WT) and G20 (HD). **B-C.** The IGV gene tracks show the distribution of active histone marks over ENCODE candidate cis-regulatory elements (cCRE) and CpG islands (CpGI), at synaptic gene promoters on chromosome 19 or at the *HOXB* cluster in WT and HD hGPCs. **D.** HD-downregulated genes *SASH1* and *SOX6* (see also **Figure S1E**) contain sites with significant H3K9ac loss at their promoters. **E.** *DiffBind* analysis found significant H3K4me3 depletion in HD (G20) compared to WT (G19) hGPCs at glial genes, which were consistently suppressed in HD across cell lines and analyses (*edgeR* and *DESeq2*) in previously reported RNA-seq data. Transformation (sign(Log_2_FC)•Log2(|Log_2_FC|+1)) was used to reduce outlier effects while maintaining directionality. Asterisks mark statistical significance; *P < 0.05; **P < 0.01; ***P < 0.001.

**Figure S2**

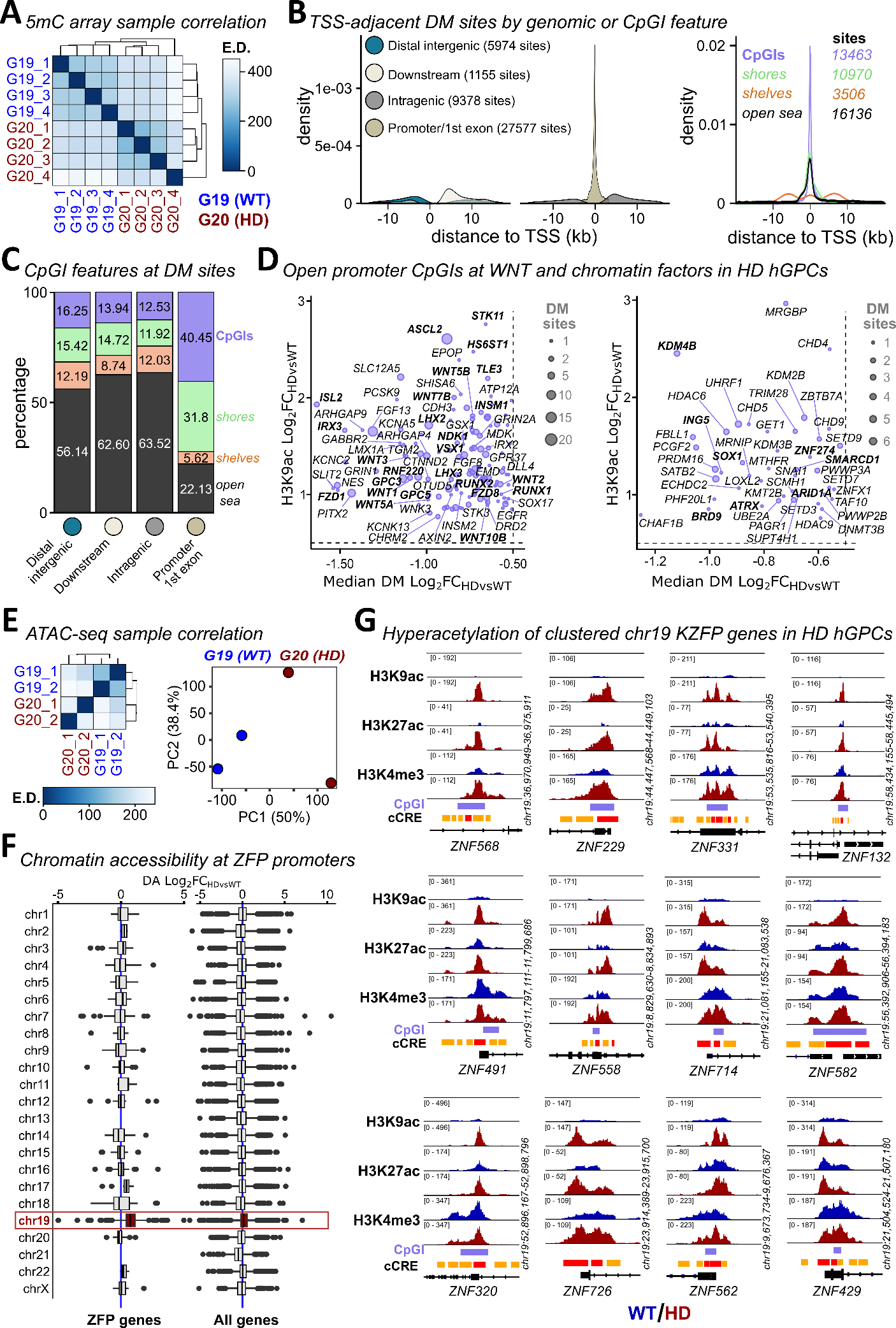

**Figure S2. Aberrantly open chromatin and DNA hypomethylation at KZFP gene promoters across chr19, related to Figure 2.**

**A.** Euclidean distance (E.D.) correlation heatmap of 5-methylcytosine (5mC) profiles in G19 (WT) vs G20 (HD) hGPCs. **B.** The distribution of differentially methylated (DM) sites (|Log_2_FC|>0.5, p<0.05) across gene TSSs are shown for annotated genomic features (left) or intersecting CpG islands (CpGIs), shores and shelves, or open sea regions (right). **C.** The proportion of significant DM sites associated with CpG islands (CpGIs), shores and shelves, or open sea regions is shown for annotated genomic features. **D.** Changes at promoter-associated sites exhibiting both significant DNA hypomethylation (Log_2_FC<-0.5 & p<0.05) and H3K9 hyperacetylation (Log_2_FC>0.5 & p<0.05) are shown for the top 10 GO terms (related to **Figure 2B**) not associated with regulation of transcription (terms #2, #5, #7, #14 in **Table S1**). DNA demethylation and H3K9 hyperacetylation were also found at chromatin remodelers (genes in terms related to “chromatin” or “histone”, were plotted alongside curated factors). **E.** Euclidean distance (E.D.) correlation heatmap and PCA of ATAC-seq samples show distinct chromatin accessibility profiles between HD and WT hGPCs. **F.** All differences in chromatin accessibility are compared per chromosome for zinc finger protein (ZFP) gene promoters versus promoters from all other genes, with chromosome 19 (chr19) highlighted in red.
**G.** The gene tracks present the accumulation of active histone marks H3K9ac, H3K27ac, and H3K4me3 in WT and HD hGPCs, across KZFP gene promoters from several chr19 clusters. Several chr19 KZFP gene promoters display H3K9ac hyperacetylation with concomitant H3K4me3/H3K27ac increases suggesting transcriptional activation.

**Figure S3**

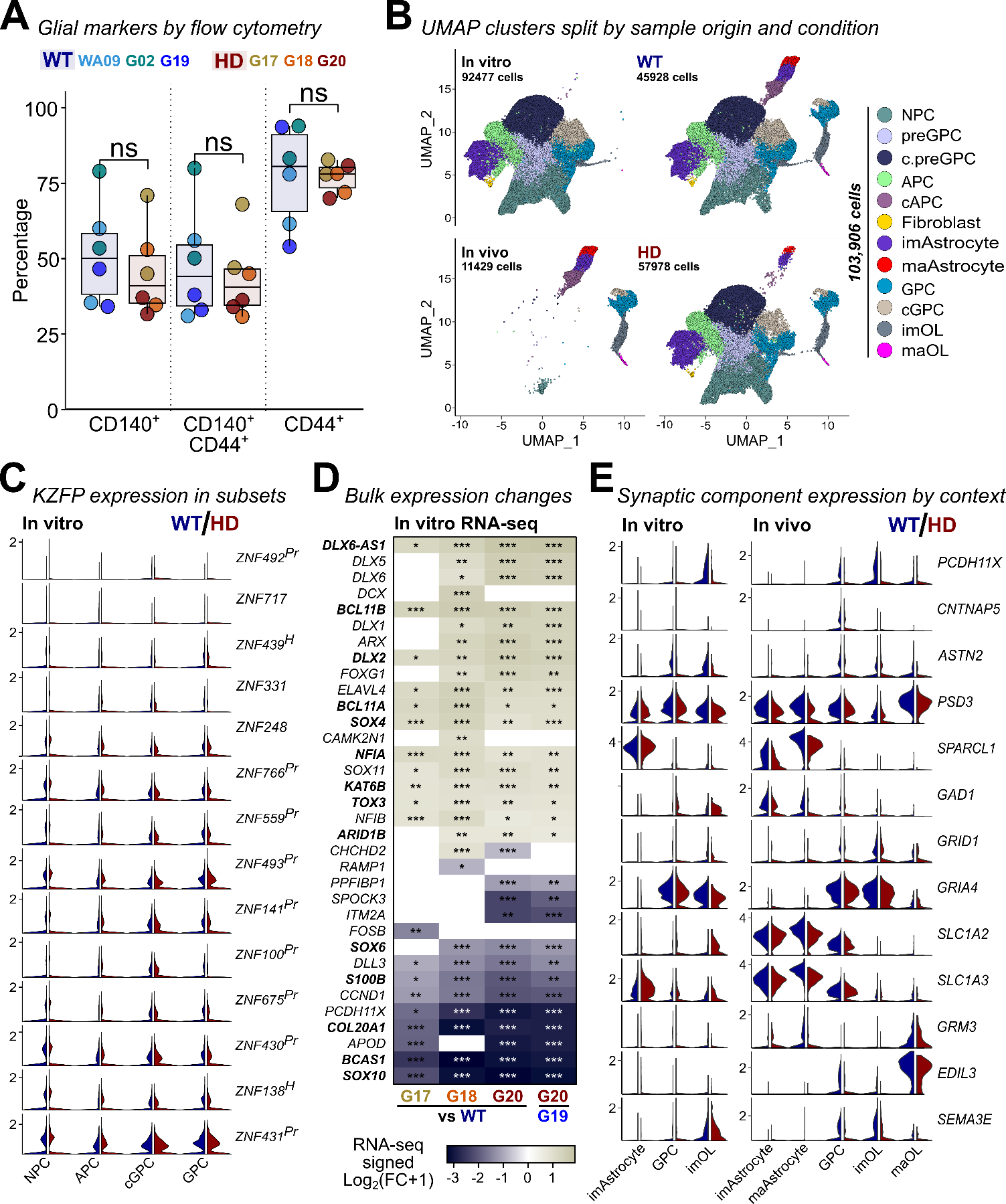

**Figure S3. Single cell transcriptomics reveal context-dependent malfunction of HD hGPCs, related to Figures 4 and 5.**

**A.** Glial inductions for scRNA-seq assessed by flow cytometry showed similar expression of glial surface markers. **B.** Uniform Manifold Approximation and Projection (UMAP) visualization of integrated in vitro hGPC and in vivo hGPC datasets split by sample origin (in vitro or in vivo) and condition (WT or HD).
**C.** The violin plots compare expression patterns for more or less abundant KZFPs across clusters in WT and HD hGPC samples in vitro. KZFPs have been marked for primate- and hominid-specificity with *Pr* and *H*, respectively.  **D.** The scRNA-seq of in vitro HD GPCs reproduce several observations from bulk RNA-seq (compare to **Figure 5A**), highlighting their developmental impairment. Signed Log_2_FC values (bulk RNA-seq) have been transformed to reduce the effect of outliers while maintaining directionality (sign(Log_2_FC)•Log2(|Log_2_FC|+1)). Asterisks mark statistical significance; *P < 0.05; **P < 0.01; ***P < 0.001. **E.** The violin plots highlight the suppression of diverse functional components across HD GPC-derived glia.

**Figure S4**

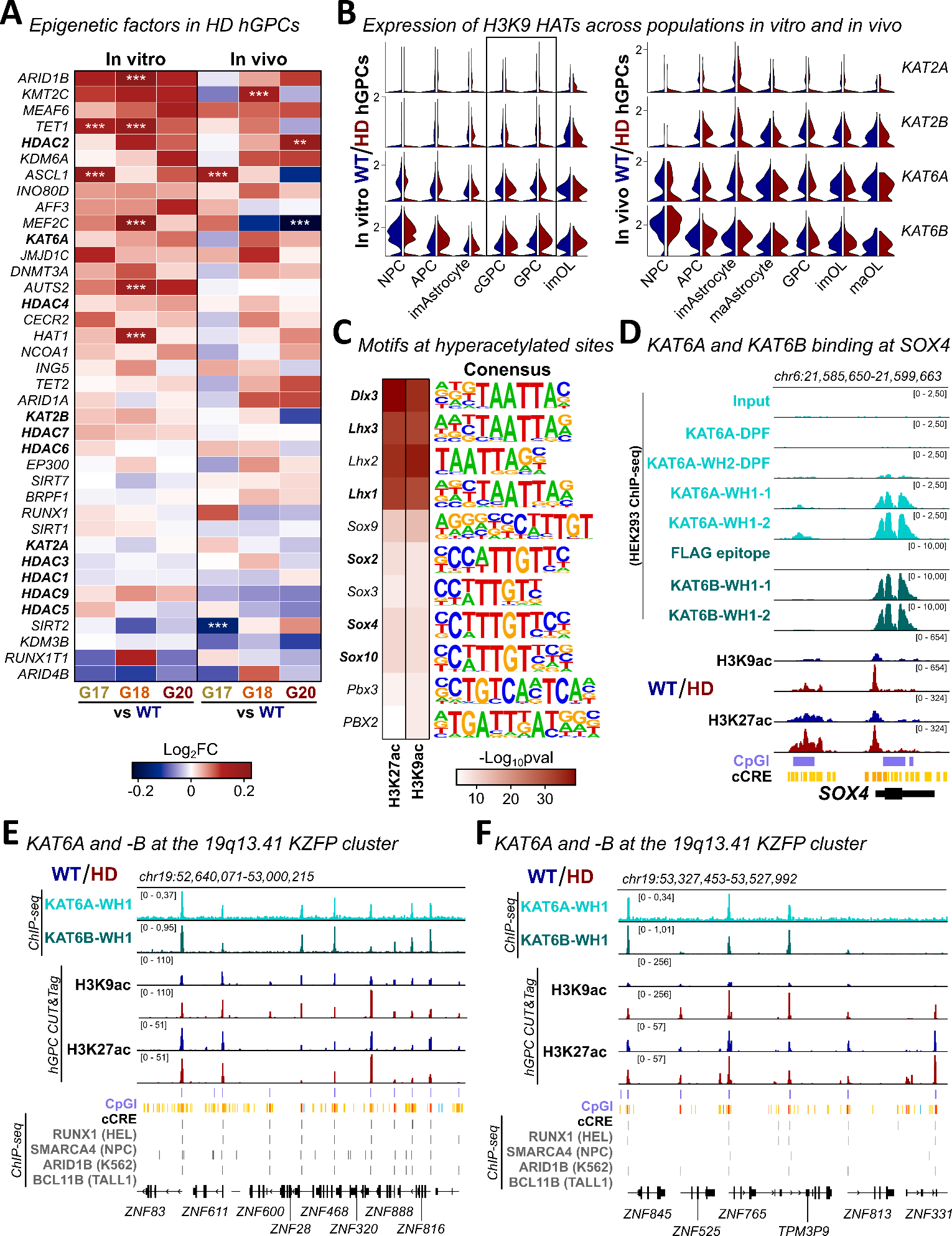

**Figure S4. MYST HATs KAT6A and KAT6B target CpGIs at chr19 KZFP gene clusters alongside key TFs, related to Figure 5.**

**A.** A variety of epigenetic regulators, such as the MYST HAT co-factors *BRPF1, ING5,* and *MEAF6,* as well as several HDACs showed only minor increases in activity in HD GPCs. Asterisks mark significance; *P < 0.05; **P < 0.01; ***P < 0.001. **B.** The expression differences for HATs *KAT2A, KAT2B, KAT6A,* and *KAT6B* in WT and HD subpopulations are compared between mature glia and NPCs from WT and HD hGPCs. **C.** Homer motif enrichment analysis carried out using the significantly hyperacetylated sites (H3K27ac and H3K9ac), showed enrichment of several SOX family TFs as well as DLX-related motifs, as illustrated by the motifs on the right. The full list of motifs can be found in **Table S2**. **D.** The gene tracks compare the KAT6A and KAT6B peaks generated from ChIP-seq of HEK293 cells following overexpression of the winged helix domain 1 (WH1, CpG-binding), WH2, or the double PHD finger domain (DPF) to their respective controls at the *SOX4* locus. **E-F.** KAT6A and KAT6B binding sites are shown across recently expanded KZFPs clusters on chr19, intersecting with recruitment sites of subunits belonging to SWI/SNF remodeling complexes. These data were sourced from public ChIP-seq datasets (listed in the methods section).

**Figure S5**

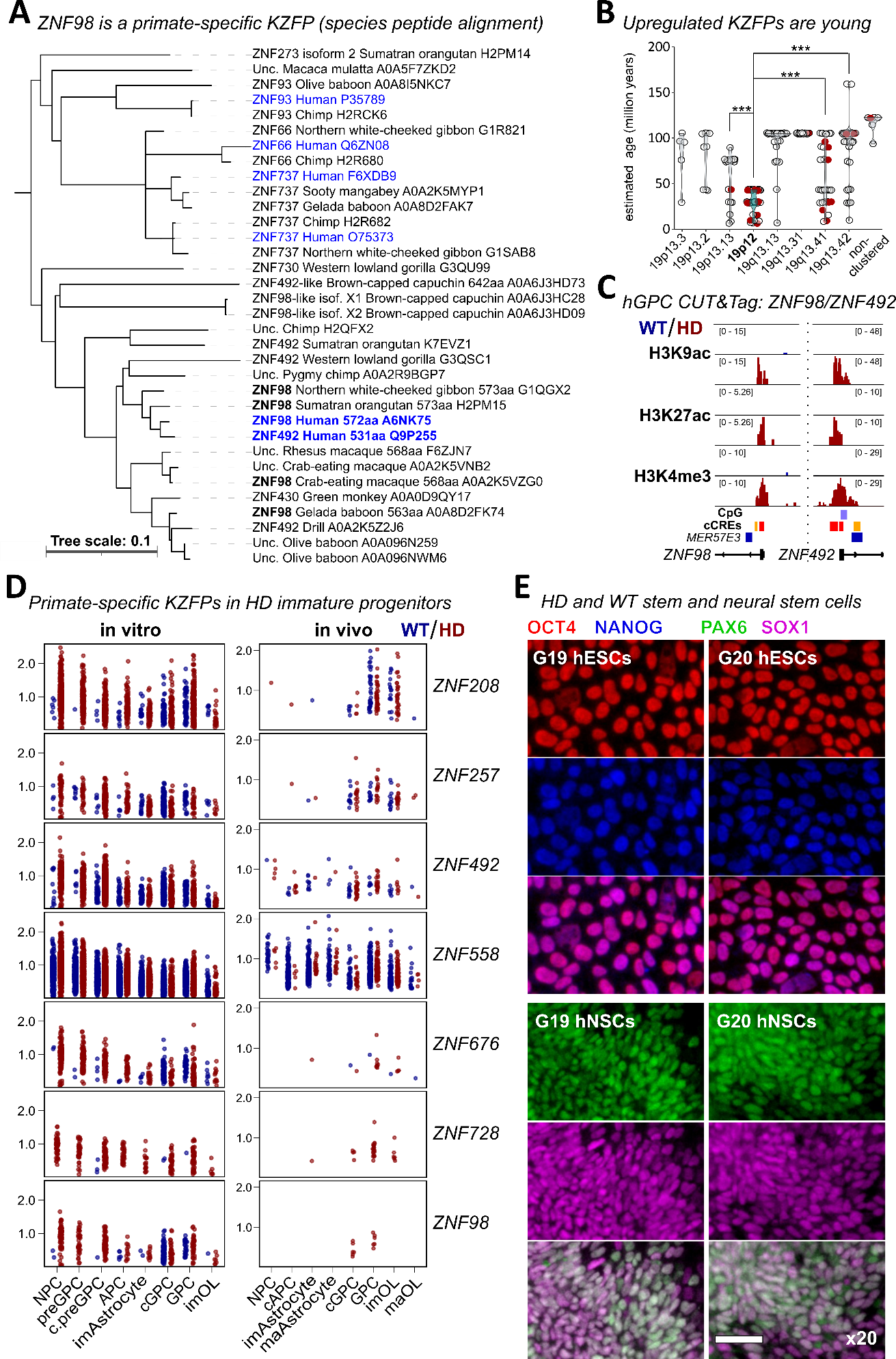

**Figure S5. The primate-specific ZNF98 exhibits enriched expression in immature populations, related to Figure 6.**

**A.** The phylogenetic tree shows the closest peptides to human ZNF98 and their species of origin identified by Uniprot blast and compared by Clustal Omega alignment. Human KZFPs are marked in blue while ZNF98 sequences from other primates are shown in bold. The 43 closest hits of 249 total are shown. The tree was constructed using iTOL. Asterisks: t-test p-values (*p<0.05; **p<0.01; ***p<0.001). **B.** Several upregulated KZFPs (G20 vs G19 comparison, related to **Figure 3D**) originate from cluster 19p12 and are among the youngest on chr19 **C.** Active chromatin marks from G19 (WT) and G20 (HD) hGPCs are presented at the *ZNF98* and *ZNF492* promoters. **D.** The expression of primate-specific KZFPs from cluster 19p12 and ZNF558 is compared across HD and WT hGPCs from in vitro and in vivo scRNA-seq datasets. **E.** Expression of pluripotency markers in hESCs and neural markers following neural induction (DIV30) was assessed by immunocytochemistry.

**Figure S6**

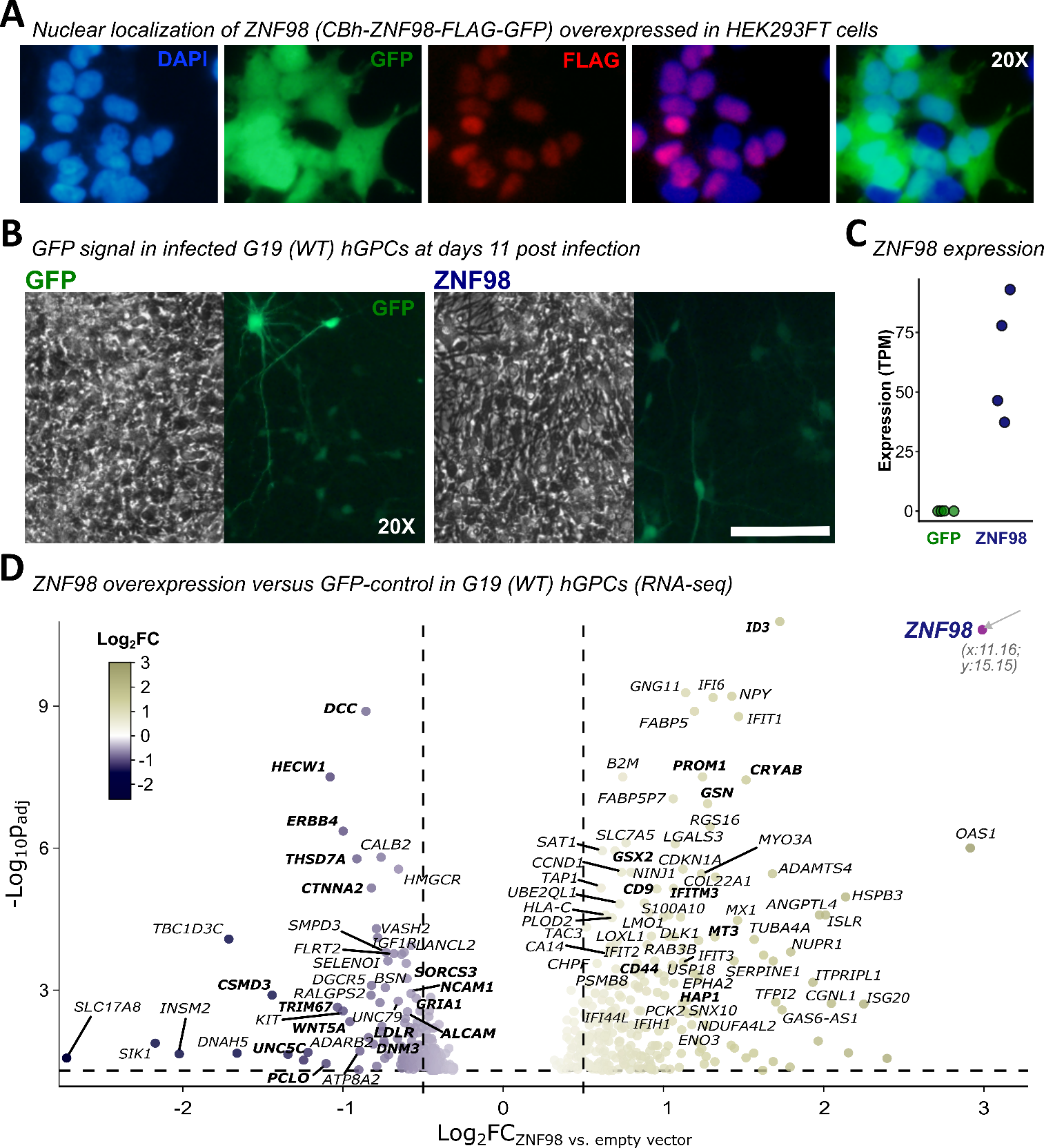

**Figure S6. ZNF98 overexpression in WT hGPCs reproduces HD glial transcriptional suppression, related to Figure 6.**

**A.** Nuclear localization of ZNF98 was confirmed by FLAG-staining in stably transduced HEK293 cells following 10 days post lentiviral transduction. **B.** ZNF98-infected G19 (WT) hGPCs were assessed 24h pre FACS isolation. **C.** RNA-seq confirms efficient ZNF98 overexpression in comparison to the hGPCs expressing the GFP control vector. **D.** Volcano plot of ZNF98-induced expression changes for genes passing the significance threshold (|Log_2_FC|>0.5, p_adj_<0.05).

**SUPPLEMENTAL TABLES**

**Table S1. Top 25 enriched GO terms for activated promoters in HD hGPCs, related to Figure 2.**

The table lists the top 25 enriched GO terms for genes with promoters exhibiting both significant DNA hypomethylation and H3K9 hyperacetylation.

| # | Term | Overlap | pval | OR |
| --- | --- | --- | --- | --- |
| 1 | Regulation Of Transcription By RNA Polymerase II (GO:0006357) | 255/2028 | 2.47E-09 | 1.56 |
| 2 | Neuron Differentiation (GO:0030182) | 38/173 | 1.30E-07 | 2.93 |
| 3 | Negative Regulation Of Transcription By RNA Polymerase II (GO:0000122) | 110/763 | 2.25E-07 | 1.78 |
| 4 | Negative Regulation Of DNA-templated Transcription (GO:0045892) | 138/1025 | 3.94E-07 | 1.65 |
| 5 | Generation Of Neurons (GO:0048699) | 36/172 | 9.48E-07 | 2.75 |
| 6 | Regulation Of DNA-templated Transcription (GO:0006355) | 228/1922 | 1.98E-06 | 1.44 |
| 7 | Potassium Ion Transmembrane Transport (GO:0071805) | 28/137 | 2.34E-05 | 2.66 |
| 8 | Potassium Ion Transport (GO:0006813) | 25/122 | 5.99E-05 | 2.67 |
| 9 | Regulation Of Canonical Wnt Signaling Pathway (GO:0060828) | 36/207 | 7.02E-05 | 2.18 |
| 10 | Midbrain Dopaminergic Neuron Differentiation (GO:1904948) | 6/10 | 7.46E-05 | 5.44 |
| 11 | Regulation Of Monoatomic Cation Transmembrane Transport (GO:1904062) | 12/39 | 9.32E-05 | 4.58 |
| 12 | Regulation Of Small GTPase Mediated Signal Transduction (GO:0051056) | 24/118 | 9.44E-05 | 2.64 |
| 13 | Positive Regulation Of Neural Precursor Cell Proliferation (GO:2000179) | 10/29 | 1.23E-04 | 5.43 |
| 14 | AC-Inhibiting G Protein-Coupled Receptor Signaling Pathway (GO:0007193) | 14/52 | 1.26E-04 | 3.80 |
| 15 | Regulation Of AMPA Receptor Activity (GO:2000311) | 9/24 | 1.28E-04 | 6.18 |
| 16 | Dendritic Spine Morphogenesis (GO:0060997) | 7/15 | 1.46E-04 | 9.01 |
| 17 | Chemical Synaptic Transmission (GO:0007268) | 43/273 | 1.58E-04 | 1.94 |
| 18 | Metanephros Development (GO:0001656) | 9/25 | 1.84E-04 | 5.80 |
| 19 | Regulation Of Intracellular Signal Transduction (GO:1902531) | 45/297 | 2.77E-04 | 1.85 |
| 20 | Positive Regulation Of Developmental Growth (GO:0048639) | 13/50 | 3.17E-04 | 3.62 |
| 21 | Extracellular Matrix Assembly (GO:0085029) | 7/17 | 3.76E-04 | 7.21 |
| 22 | Heterochromatin Organization (GO:0070828) | 11/39 | 4.24E-04 | 4.05 |
| 23 | Canonical Wnt Signaling Pathway (GO:0060070) | 15/65 | 4.62E-04 | 3.10 |
| 24 | Negative Regulation Of Cell Adhesion (GO:0007162) | 16/72 | 4.76E-04 | 2.95 |
| 25 | Neurotransmitter Transport (GO:0006836) | 13/52 | 4.80E-04 | 3.44 |

**Table S2. Aberrant availability at chr19 KZFP genes in HD hGPCs. Standalone excel file, related to Figures 1, 2 and S4.**

The standalone Table_S2.xlsx (Excel) lists those chromosome 19 ZFP transcripts which exhibit increased accessibility (ATAC-seq) in HD (G20) relative to WT (G19) hGPCs. It also separately lists all ZFP transcripts that are significantly dysregulated (RNA-seq) in HD compared to WT hGPCs (related to **Figures 2** and **3**, respectively). KZFPs previously identified as primate (Pr) or hominid-specific (H) are marked accordingly.

**Table S3. Transcriptional analyses of HD hGPCs. Standalone excel file, related to Figures 5, 6, S4 and S6.**

The standalone Table_S3.xlsx (Excel) contains all motifs identified by *RcisTarget*.

**Table S4. ZNF98 and ZNF492 peptide alignment**

The high similarity between the amino acid sequences of ZNF98 and ZNF492 is seen in the following alignment (Clustal W 1.83 multiple sequence alignment). The C2H2 zinc finger domains are underlined, differences are shown in red, while the KRAB domains are marked in grey.

| sp\|Q9P255\|ZN492 sp\|A6NK75\|ZNF98 | -----------------------------------------MLENYRNLVFVGIAASKPD  MPGPLGSLEMGVLTFRDVALEFSLEEWQCLDTAQQNLYRNVMLENYRNLVFVGIAASKPD |
| --- | --- |
| sp\|Q9P255\|ZN492 sp\|A6NK75\|ZNF98 | LITCLEQGKEPWNVKRHEMV**A**EPPVV**C**SYFA**R**DLWPKQGKKNYFQKVILR**R**YKKCG**C**ENL LITCLEQGKEPWNVKRHEMV**T**EPPVV**Y**SYFA**Q**DLWPKQGKKNYFQKVILR**T**YKKCG**R**ENL |
| sp\|Q9P255\|ZN492 sp\|A6NK75\|ZNF98 | QLRKYCKSMDECKVHKECYNGLNQCLTTTQNKIFQ**C**DKYVKVFHKFSNSNRH**T**I**R**HTGKK QLRKYCKSMDECKVHKECYNGLNQCLTTTQNKIFQ**Y**DKYVKVFHKFSNSNRH**K**I**G**HTGKK |
| sp\|Q9P255\|ZN492 sp\|A6NK75\|ZNF98 | S**FKCKECEKSFCMLSHLAQHKRIH**SGEKP**YKCKECGKAYNETSNLSTHKRIH**TGKKP**YKC**  S**FKCKECEKSFCMLSHLAQHKRIH**SGEKP**YKCKECGKAYNEASNLSTHKRIH**TGKKP**YKC** |
| sp\|Q9P255\|ZN492 sp\|A6NK75\|ZNF98 | **EECGKAFNRLSHLTTHKIIH**TGKKP**YKCEECGKAFNQSANLTTHKRIH**TGEKP**YKCEECG**  **EECGKAFNRLSHLTTHKIIH**TGKKP**YKCEECGKAFNQSANLTTHKRIH**TGEKP**YKCEECG** |
| sp\|Q9P255\|ZN492 sp\|A6NK75\|ZNF98 | **RAFSQSSTLTAHKIIH**AGEKP**YKCEECGKAFSQSSTLTTHKIIH**TGEKF**YKCEECGKAFS**  **RAFSQSSTLTAHKIIH**AGEKP**YKCEECGKAFSQSSTLTTHKIIH**TGEKF**YKCEECGKAFS** |
| sp\|Q9P255\|ZN492 sp\|A6NK75\|ZNF98 | **QLSHLTTHKRIH**SGEKP**YKCEECGKAFKQSSTLTTHKRIH**AGEKF**YKCEVCSKAFSRFSH RLSHLTTHKRIH**SGEKP**YKCEECGKAFKQSSTLTTHKRIH**AGEKF**YKCEVCSKAFSRFSH** |
| sp\|Q9P255\|ZN492 sp\|A6NK75\|ZNF98 | **LTTHKRIH**TGEKP**YKCEECGKAFNLSSQLTTHKIIH**TGEKP**YKCEECGKAFNQSSTLSKH**  **LTTHKRIH**TGEKP**YKCEECGKAFNLSSQLTTHKIIH**TGEKP**YKCEECGKAFNQSSTLSKH** |
| sp\|Q9P255\|ZN492 sp\|A6NK75\|ZNF98 | **KVIH**TGEKP**YKYEECGKAFNQSSHLTTHKMIH**TGEKP**YKCEECGKAFNNSSILNRHKMIH KVIH**TGEKP**YKCEECGKAFNQSSHLTTHKMIH**TGEKP**YKCEECGKAFNNSSILNRHKMIH** |
| sp\|Q9P255\|ZN492 sp\|A6NK75\|ZNF98 | TGEKLYKPESCNNACDNIAKISKYKRNCAGEK  TGEKLYKPESCNNACDNIAKISKYKRNCAGEK |
